# Spatiotemporal distribution of the glycoprotein pherophorin II reveals stochastic geometry of the growing ECM of *Volvox carteri*

**DOI:** 10.1101/2024.12.06.625376

**Authors:** Benjamin von der Heyde, Anand Srinivasan, Sumit Kumar Birwa, Eva Laura von der Heyde, Steph S.M.H. Höhn, Raymond E. Goldstein, Armin Hallmann

**Author notes:** Joint first author (Benjamin von der Heyde, Anand Srinivasan).

## Abstract

The evolution of multicellularity involved the transformation of a simple cell wall of unicellular ancestors into a complex, multifunctional extracellular matrix (ECM). A suitable model organism to study the formation and expansion of an ECM during ontogenesis is the multicellular green alga *Volvox carteri*, which, along with the related volvocine algae, produces a complex, self-organized ECM composed of multiple substructures. These self-assembled ECMs primarily consist of hydroxyproline-rich glycoproteins, a major component of which is pherophorins. To investigate the geometry of the growing ECM, we fused the *yfp* gene with the gene for pherophorin II (PhII) in *V. carteri*. Confocal microscopy reveals PhII:YFP localization at key structures within the ECM, including the boundaries of compartments surrounding each somatic cell and the outer surface of the organism. Image analysis during the life cycle allows the stochastic geometry of those growing compartments to be quantified. We find that their areas and aspect ratios exhibit robust gamma distributions and exhibit a transition from a tight polygonal to a looser acircular packing geometry with stable eccentricity over time, evoking parallels and distinctions with the behavior of hydrated foams. These results provide a quantitative benchmark for addressing a general, open question in biology: How do cells produce structures external to themselves in a robust and accurate manner?

---

Throughout the history of life, one of the most significant evolutionary transitions was the formation of multicellular eukaryotes. In most lineages that evolved multicellularity, including animals, fungi and plants, the extracellular matrix (ECM) has been a key mediator of this transition by connecting, positioning, and shielding cells [1–4]. The same holds for multicellular algae such as *Volvox carteri* (Chlorophyta) and its multicellular relatives within volvocine green algae which developed a remarkable array of advanced traits in a comparatively short amount of evolutionary time—oogamy, asymmetric cell division, germ-soma division of labor, embryonic morphogenesis and a complex ECM [5–8]—rendering them uniquely suited model systems for examining evolution from a unicellular progenitor to multicellular organisms with different cell types [5, 6, 8–10]. In particular, *V*.*carteri* ‘s distinct and multilayered ECM makes it a model organism for investigating the mechanisms underlying ECM growth and its effects on the positioning of the cells that secrete its components. Building on recently established protocols for stable expression of fluorescently labeled proteins in *V. carteri* [11–13] we present here a new transgenic strain revealing localization of the glycoprotein pherophorin II and the first *in vivo* study of the stochastic geometry of a growing ECM.

*V. carteri* usually reproduces asexually (Fig. 1A). Sexual development is triggered by exposure to heat or a species-specific glycoprotein sex inducer, which results in development of sperm-packet-bearing males and egg-bearing females [14–17]. In the usual asexual development, a *V. carteri* organism consists of ∽2000 biflagellate somatic cells resembling *Chlamydomonas* in their morphology, arranged in a monolayer at the surface of a sphere and approximately 16 much larger, nonmotile, asexual reproductive cells (gonidia) that constitute the germline, lying just below the somatic cell layer [5– 7, 18, 19]. The somatic cells are specialized for ECM biosynthesis, photoreception and motility; for phototaxis, they must be positioned correctly within the ECM at the surface of the organism [14, 20, 21].

**FIG. 1.**
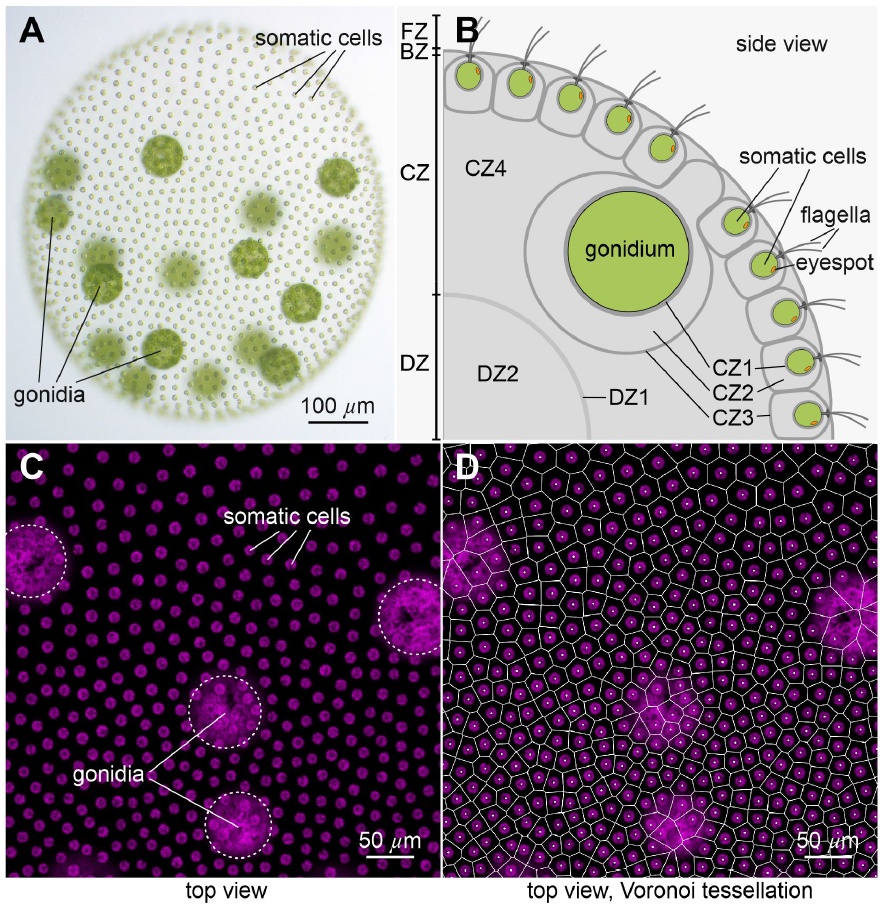
Phenotype, ECM architecture, and cell distribution of *V. carteri*. (A) Adult female (wild-type). Somatic cells each bear an eyespot and two flagella, while deeper-lying gonidia are flagella-less. All cells have a single large chlorophyll-containing chloroplast (green). (B) Zones of the ECM, as described in text. (C) Image utilizing autofluorescence of the chlorophyll (magenta) to define cell positions. (D) Semiautomated readout of somatic cell positions in (C) followed by a Voronoi tessellation gives an estimate of the ECM neighborhoods of somatic cells.

The ECM of *V. carteri* has been studied in the past decades from the perspective of structure and composition, developmental, mechanical properties, cellular interactions, molecular biology and genetics, and evolution [11, 14, 20–22]. In the ontogenesis of *V. carteri*, ECM biosynthesis in juveniles, inside their mothers, begins only after all cell divisions and the process of embryonic inversion are completed. Cells of the juveniles then continuously excrete large quantities of ECM building blocks which are integrated into their ECM, producing an enormous increase in the size of the juveniles; within 48 hours the volume increases almost 3000-fold as the diameter increases from ∽ 70 *µ*m to ∽ 1 mm, raising the question of how relative cell positions are affected by growth. In the adult organism, the ECM accounts for up to 99% of the organism’s volume and consists of morphologically distinct structures with a defined spatial arrangement.

Based on electron microscopy, a nomenclature was established [20] that defines four main ECM zones: flagellar zone (FZ), boundary zone (BZ), cellular zone (CZ) and deep zone (DZ), which are further subdivided (Fig. 1B). The CZ3 forms the ECM compartment boundaries of individual cells and the BZ constitutes the outer surface of the organism. CZ3 and BZ show a higher electron density than the ECM within the compartments or the ECM in the interior below the cell layer (CZ4 and DZ2) [6, 7, 20]. CZ3 and BZ therefore appear to consist of a firmer, more robust ECM, while CZ4 and DZ2 appear to be more gelatinous.

The ECM of volvocine algae predominantly consists of hydroxyproline-rich glycoproteins (HRGPs), which are also a major component of the ECM of embryophytic land plants [14, 21, 23–25], but contains no cellulose. In *V. carteri* and other volvocine algae, a large family of HRGPs, the pherophorins, are expected to constitute the main building material of the ECM [17, 22, 26–29]. Just in *V. carteri*, 118 members of the pherophorin family were discovered in the genome [11, 30]; pherophorin genes are typically expressed in a cell type-specific manner [31] that can be constitutive or induced either by the sex inducer or wounding [17, 22, 26, 28, 29, 32]. Pherophorins have a dumbbell-like structure: two globular domains separated by a rod-shaped, highly proline-rich domain that varies considerably in length [14, 21, 28, 29]. These prolines undergo post-translational modification to hydroxyprolines; pherophorins are also strongly glycosylated [14, 21, 22, 33, 34]. For two pherophorins the polymerization into an insoluble fibrous network was shown in vitro [26]; some have already been localized in the ECM or in ECM fractions [11, 13, 22, 27, 29].

To investigate the ECM, we determined that pherophorin II (PhII) was the appropriate one to label fluorescently, as it is firmly integrated into the ECM. It is a 70 kDa glycoprotein with strong and immediate inducibility by the sex inducer [17, 35, 36]. Because it can only be extracted under harsh conditions, it is thought to be a component of the insoluble part of the somatic CZ [17, 35, 36]. We fused one of the nine [28, 30] gene copies of PhII with the *yfp* gene. The corresponding DNA construct was stably integrated into the genome of *V. carteri* transformants by particle bombardment. The expression of fluorescent PhII:YFP was confirmed in transgenic mutants through CLSM, where it was found in the compartment boundaries of the cellular and boundary zones. This allows the proportion of ECM secreted by individual somatic cells to be determined along with the stochastic geometry of these ECM structures during development.

We suggest that a detailed study of the geometry of ECM structures during growth in *Volvox* made possible by this strain will provide insight into a more general question in biology: How do cells robustly produce structures external to themselves? There is no physical picture of how the intricate geometry of the *Volvox* ECM arises through what must be a self-organized process of polymer crosslinking [37]. While information on a structure’s growth dynamics can be inferred from its evolving shape, as done for animal epithelial cells [38], this connection has not been made for *Volvox*.

Recently [39], the somatic cell locations of *V. carteri* were determined using their chlorophyll autofluorescence, from which the ECM neighborhoods were determined by 2D Voronoi tessellation, as in Figs. 1C,D. While the somatic cells appear to be arranged in a quasi-regular pattern, this analysis reveals that their neighborhood areas exhibited a broad, skewed k-gamma distribution. Subsequent work has shown that such distributions may arise from bursty ECM production at the single-cell level [40]. While the nearest-space-allocation rule assumed by Voronoi tessellations is a reasonable first approximation of the geometry of the somatic CZ3, the validity of this approximation remains unclear. To investigate this and to answer the more general questions raised above, we performed geometric analyses of the structures illuminated by localized PhII:YFP, using metrics (Table 1) which give insight into the coexistence of local geometric stochasticity and global robustness during growth. In particular, we draw parallels between the temporal evolution of ECM structures and wet foams.

**TABLE 1.**
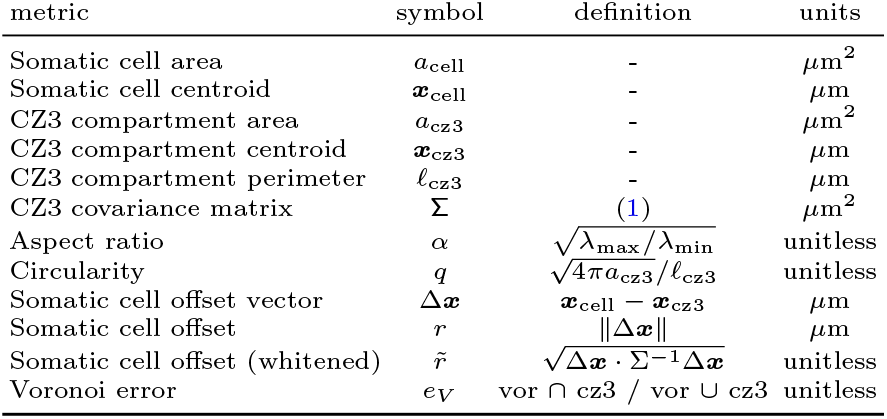
Metric definitions.

## RESULTS

### A. Vector construction and generation of transformants expressing PhII:YFP

The pherophorin II gene (*phII*) was cloned from *V. carteri* genomic DNA including its promoter, 5’ and 3’ UTRs and all seven introns (Fig. 2). A sequence coding for a flexible penta-glycine spacer and the codon-adapted coding sequence of *yfp* were inserted directly upstream of the *phII* stop codon to produce a *phII-yfp* gene fusion. The obtained vector pPhII-YFP (Fig. 2) was sequenced and then used for stable nuclear transformation of the nitrate reductase-deficient *V. carteri* recipient strain TNit-1013 by particle bombardment. To allow for selection, the non-selectable vector pPhII-YFP was cotransformed with the selectable marker vector pVcNR15, which carries an intact *V. carteri* nitrate reductase gene and, thus, complements the nitrate reductase deficiency of strain TNit-1013. Screening for transformants was then achieved by using medium with nitrate as nitrogen source. The transformants were investigated for stable genomic integration of the vector and, via confocal microscopy, for expression of the fluorescent protein at sufficient levels throughout their life cycle.

**FIG. 2.**
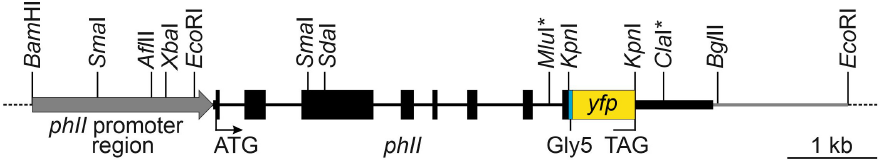
Schematic diagram of the transformation vector pPhII-YFP. The vector carries the complete *phII* gene including its promoter region. Exons (introns) are shown as black boxes (thin lines). Directly upstream of the TAG stop codon, a 0.7 kb fragment coding for a flexible penta-glycine spacer (cyan) and the codon-adapted coding sequence of *yfp* (mVenus) (yellow) were inserted in frame using an artificially introduced *Kpn*I site. The *Kpn*I was inserted before into the section between *Mlu*I and *Cla*I (asterisks). The vector backbone (dashed lines) comes from pUC18. For details, see Methods; the sequence of the vector insert is in SI Appendix, Fig. S3.

### B. In vivo localization of pherophorin II

As expected and detailed below, there are continuous changes in the amount of ECM expansion over the life cycle. While we primarily examine the parental somatic cell layer and surrounding ECM, the developmental stage of the next generation is used for a precise definition of five key stages which we focused on in the comparative analyses (Fig. 3). Stage I: Freshly hatched young adults of equivalent circular radius (defined in SI Appendix, §2.B) *R* = 106 ± 6 *µ*m, containing immature gonidia. Stage II (∽ 15 h after hatching): Middle-aged adults with *R* = 221 ±22 *µ*m containing early embryos (4-8 cell stage). Stage III (∽21 h after hatching): Older middle-aged adults with *R* = 244± 15 *µ*m containing embryos before inversion. Stage IV (∽ 36 h after hatching): Old adults of radius *R* = 422 ±6 *µ*m containing fully developed juveniles. Stage S: Sexually developed adult females of radius *R* = 265 ±29 *µ*m bearing egg cells. Since expression of PhII is induced by the sex-inducer protein [17, 35, 36], in stages I to IV with vegetative phenotype, the sex-inducer protein was added 24 hours before microscopy to increase PhII expression; after such a short incubation with the sex inducer, the females still show an unchanged cleavage program and the vegetative phenotype. To obtain a changed cleavage program and a fully developed sexual phenotype (S), the females were sexually induced 72 hours before microscopy.

**FIG. 3.**
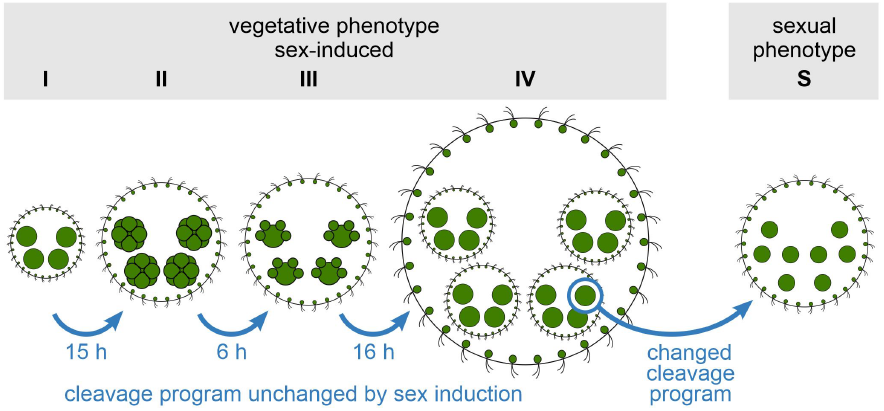
Stages of development of *V. carteri* females. Localization of PhII and quantification of ECM features were studied in 5 stages, four with vegetative (asexual) phenotype (I-IV) and one with the fully developed sexual phenotype (S). In stages I-IV, incubation with the sex inducer was short enough to increase PhII expression for adequate localization without changing the vegetative cleavage program.

### C. Pherophorin II is localized in the compartment borders of individual cells

As expected, PhII:YFP is only found in the extracellular space within the ECM. It is detectable at all developmental stages after embryonic inversion, which marks the beginning of ECM biosynthesis [34, 41], and in the ECM of organisms with the phenotypes of both vegetative and sexual development. At a first glance, in a top view, PhII:YFP appears to form a polygonal pattern at the surface of each post-embryonic spheroid, with a single somatic cell near the center of each compartment (Fig. 4). PhII:YFP is also found in the ECM compartment boundaries (CZ3) of the gonidia, which are located below the somatic cell layer (Figs. 1B and 4). These observations hold at all stages after embryonic inversion, even if the shape of the compartment boundaries varies. On closer inspection (Fig. 5), it becomes apparent that: i) the shapes of the somatic ECM compartment boundaries include a mix of hexagons, heptagons, pentagons and other polygons, as well as circles and ovals, ii) the angularity of the compartment boundaries changes in the course of expansion of the organism during development, i.e. the compartments become increasingly circular (less polygonal), and iii) each cell builds its own ECM compartment boundary. The observed localization of PhII is shown schematically in Fig. 12.

**FIG. 4.**
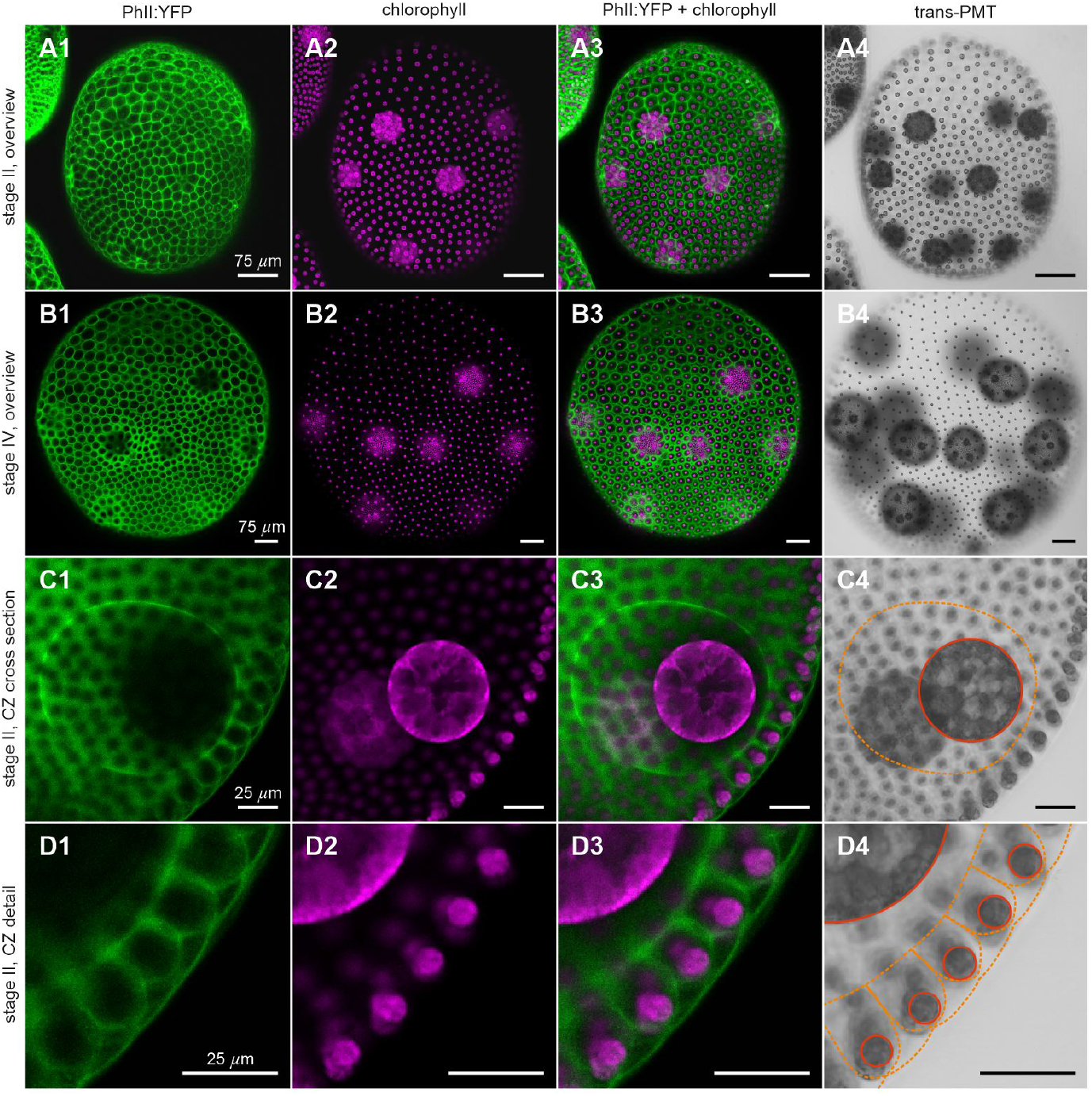
Localization of PhII:YFP in whole, middle-aged (early stage II) and old adults (stage IV) in top view and magnified cross section through CZ. Sexually induced transformants expressing the *phII* :*yfp* gene under the control of the endogenous *phII* promoter were analyzed in vivo for the localization of the PHII:YFP fusion protein. PhII:YFP is located in the CZ3 of both somatic cells and gonidia as well as in the BZ. Top views of whole organism: (A) stage II immediately before the first cleavage of the gonidia inside; and (B) stage IV) before hatching of the fully developed juveniles. (C) Cross section through CZ in early stage II. (D4) Magnified view of the outer most ECM region. Column 1: YFP fluorescence of the PhII:YFP protein (green). Column 2: Chlorophyll fluorescence (magenta). Column 3: Overlay of YFP and chlorophyll fluorescence. Column 4: Transmission-PMT (trans-PMT). PhII:YFP-stained ECM boundaries are highlighted in orange and cell boundaries in red.

**FIG. 5.**
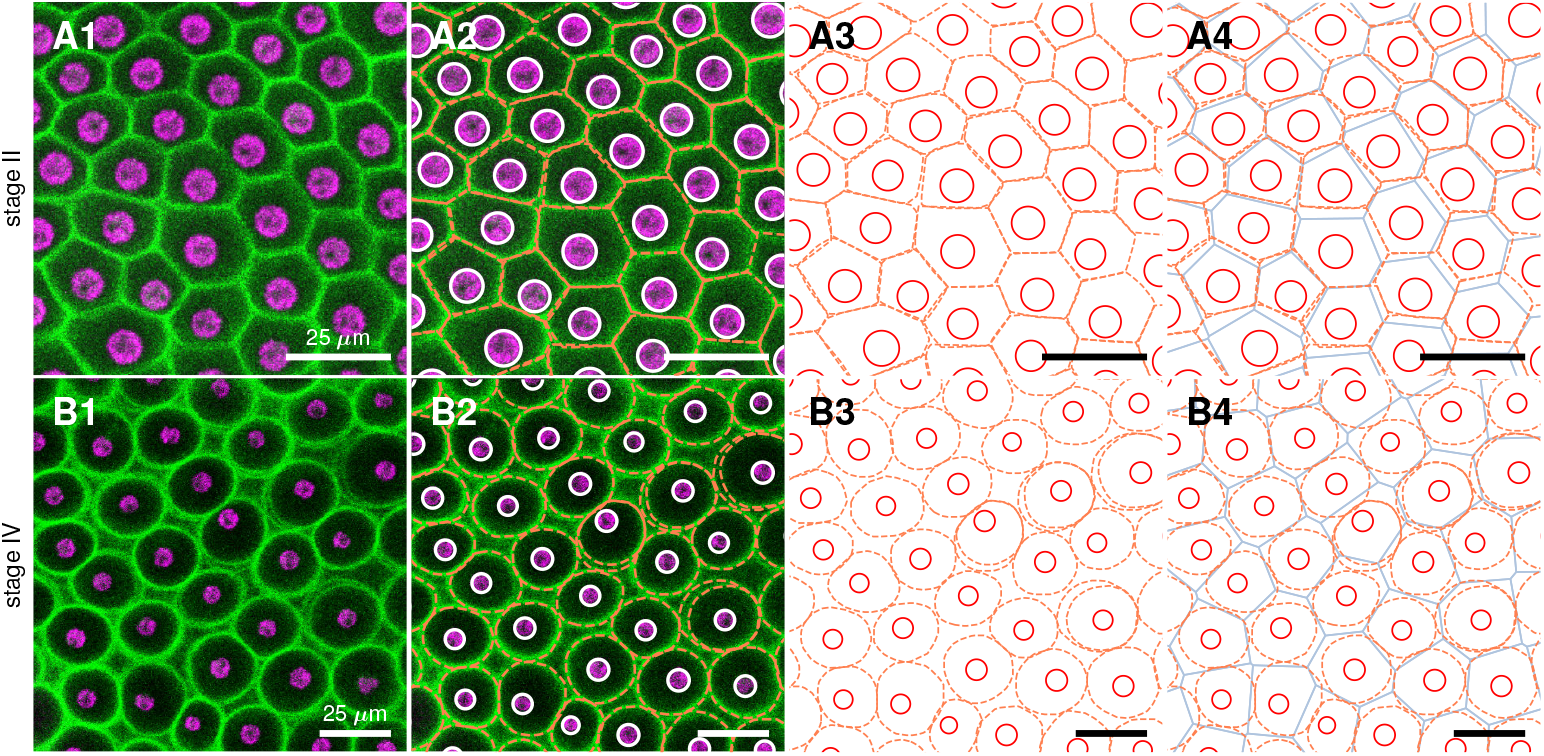
As in Fig. 4, but a close-up of PhII:YFP localization in top view. (A) Stage II. Magnified view of the somatic CZ3 compartments (1), identified by orange in 2 along with cell boundaries (white), shown together in 3 and in 4 with underlaid Voronoi tessellation (blue) and cell boundaries in red. (B) In stage IV the CZ3 compartments become more bubble shaped and individual CZ3 walls separate from the neighbors, leaving extracompartmental ECM space. Double-walls also appear, highlighted in B3.

Especially in the early stages I and II of development, the fluorescent ECM compartment borders are predominantly pentagonal or hexagonal (Figs. 4A and 5A) with a rough ratio of 1:2 (SI Appendix, Fig. S4). By stage IV they become more rounded and the interstices we term “extracompartmental ECM spaces” between the compartments increase in size and number (Figs. 4B and 5B). Since the compartment boundaries of adjacent cells are close to each other in early stages, the two compartments appear to be separated by a single wall. In later stages, when the boundaries are more circular, it becomes apparent that it is a double wall; thus, each somatic cell produces its own boundary (Figs. 5A,B).

As shown, PhII:YFP is a component of the CZ3 of both somatic cells and gonidia (Figs. 1B, 4 and 5). It seems to be a firmly anchored building block there, as the observed structure is highly fluorescent and yet sharply demarcated from other non-fluorescent adjacent ECM structures. If the PhII:YFP protein was prone to diffusion, one would expect a brightness gradient starting from the structures. There are no interruptions in the fluorescent labeling of boundaries associated with passages between neighboring compartments. As the boundaries are so close together in early stages, only the thickness of the double wall can be determined and then halved; we estimate stage II single wall thickness as 1.6 ± 0.4 *µ*m and the similar value 1.9 ± 0.5 *µ*m in stage IV.

A comparison of the PhII:YFP-stained structures with descriptions of ECM structures from electron microscopy shows that the PhII:YFP location corresponds exactly to that of the ECM structure CZ3 [20]. This applies to the entire period from the beginning of ECM biosynthesis after embryonic inversion to the maximally grown old adult.

The relatively regular pattern of compartments (Fig. 5A) is disturbed by the gonidia (later embryos) which, being far larger than somatic cells, are pushed under the somatic cell layer. The CZ3 compartments of the deeper gonidia nevertheless extend to the surface (Fig. 6). The surrounding somatic CZ3 compartments are elongated in the direction of the gonidial CZ3 protrusions to the surface (Fig. 6). Since the PhII:YFP-stained CZ3 structure completely encloses each cell, these walls cannot be impermeable; even ECM proteins must pass through them. One indicator for this is that the enormous growth of the spheroid during development requires the cell-free areas outside the ECM compartments of individual cells to increase immensely in volume. This applies in particular to the deep zone, but also to the areas between the compartments. All ECM material required for this increase in volume can only be produced and exported by the cells and must then pass through the ECM compartment borders of individual cells.

**FIG. 6.**
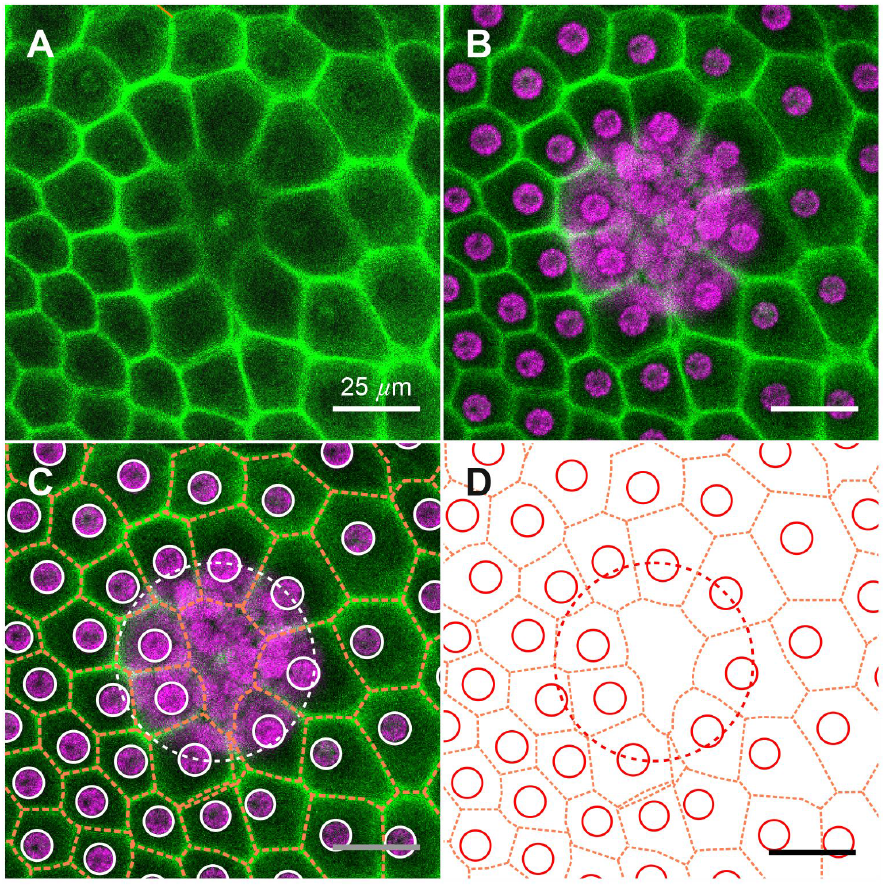
Magnified top view of CZ3 above a reproductive cell in early stage II. (A) PhII:YFP fluorescence (green). (B) overlay with chlorophyll fluorescence (magenta). (C) overlay with highlighted CZ3 (orange) and cell boundaries (white). (D) as in (C) with only cell boundaries (red) and CZ3 (orange).

### D. Quantification of somatic CZ3 geometry

The localization of PhII at the CZ3 compartment boundaries allows us to carry out the first quantitative analyses of their geometry, both along the PA axis and through the life cycle stages. A semi-automated image analysis pipeline (SI Appendix, §2) reveals geometric features described in Table 1 and Fig. 7. A total of 29 spheroids across five developmental stages were analyzed: 7 in stage I, 5 each in stages II, III, and IV of the asexual life cycle, and 7 in the sexual life cycle (stage S).

**FIG. 7.**
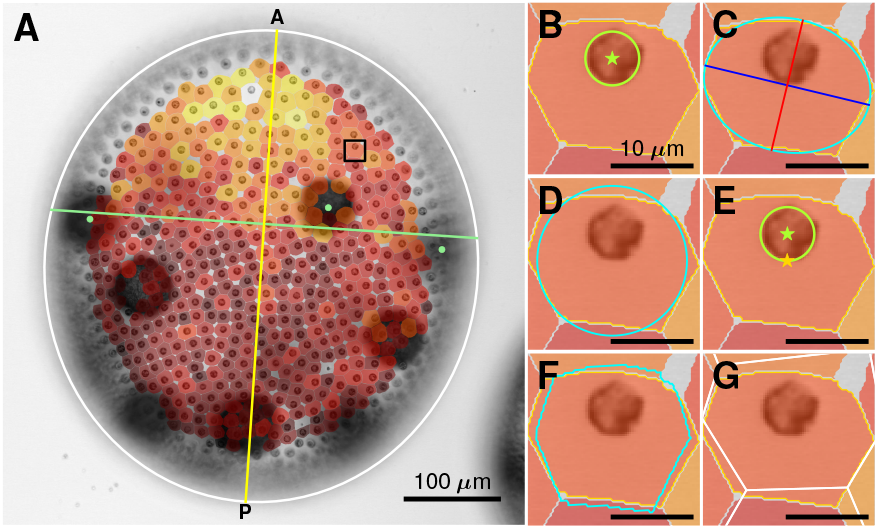
Geometric features of cell/compartments pairs. (A) Trans-PMT image of stage III spheroid, with elliptical outline (white), estimated PA axis (yellow) that is orthogonal to line through gonidia (green dots). Overlaid are segmentations of CZ3 compartments, colored dark to light by size. (B-G) Schematics of geometric features computed from cell (green) and compartment (yellow) boundaries, as indicated: (B) *a*_cell_, *a*_cz3_, (C) aspect ratio *α* and corresponding ellipse (cyan), (D) deviation from a circle of the same area (cyan), (E) offset of cell center of mass (green star) from compartment center of mass (yellow star), (F) whitening transform of the compartment (cyan), and (G) Voronoi tessellation (white) error *e*_*V*_.

The PA axis of *V. carteri* spheroids corresponds to their swimming direction, and along this axis the distance between the somatic cells and the size of their eyespots decreases toward the posterior pole (Fig. 1A). Offspring (gonidia, embryos, and daughter spheroids) are mainly located in the posterior hemisphere. We approximate the PA axis by a line passing through the center of the spheroid and normal to the best-fit line passing through manually identified juveniles in the anterior (Fig. 7A). This estimated PA axis is typically well-approximated by the elliptical major axis of the spheroid (SI Appendix, Fig. S5).

The metrics presented in Fig. 7 (see SI Appendix, §2) are derived from the compartment shape outline or from moments of area. The matrix M_2_ of second moments of area and the normalized matrix ∑ = M_2_*/a*_cz3_, with the former defined as

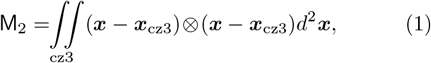

are interpretable as elastic strain tensors with respect to unit-aspect-ratio shapes. ∑’s eigenvalues *λ*_max_, *λ*_min_ (whose square roots define the major, minor axis lengths) are principal stretches of this deformation, yielding the aspect ratio and other quantities defined in Table 1. Overall, we measure changes in the moments of area (SI Appendix, §2.B) to quantify ECM geometry during growth; the 0^th^ gives the area increase, the 1^st^ quantifies migration of compartment centroids with respect to cells, the 2^nd^ gives the change in anisotropy (Table 1). The sum of second moments reveals changes in crystallinity of the entire CZ3 configuration, as shown in § G.

### E. CZ3 geometry along PA axis during the life cycle

#### 1. Anterior CZ3 compartments expand toward end of life cycle

Fig. 8A1-2 shows that the somatic cell area increases modestly, by ∽ 10%, along the PA axis at all stages.

**FIG. 8.**
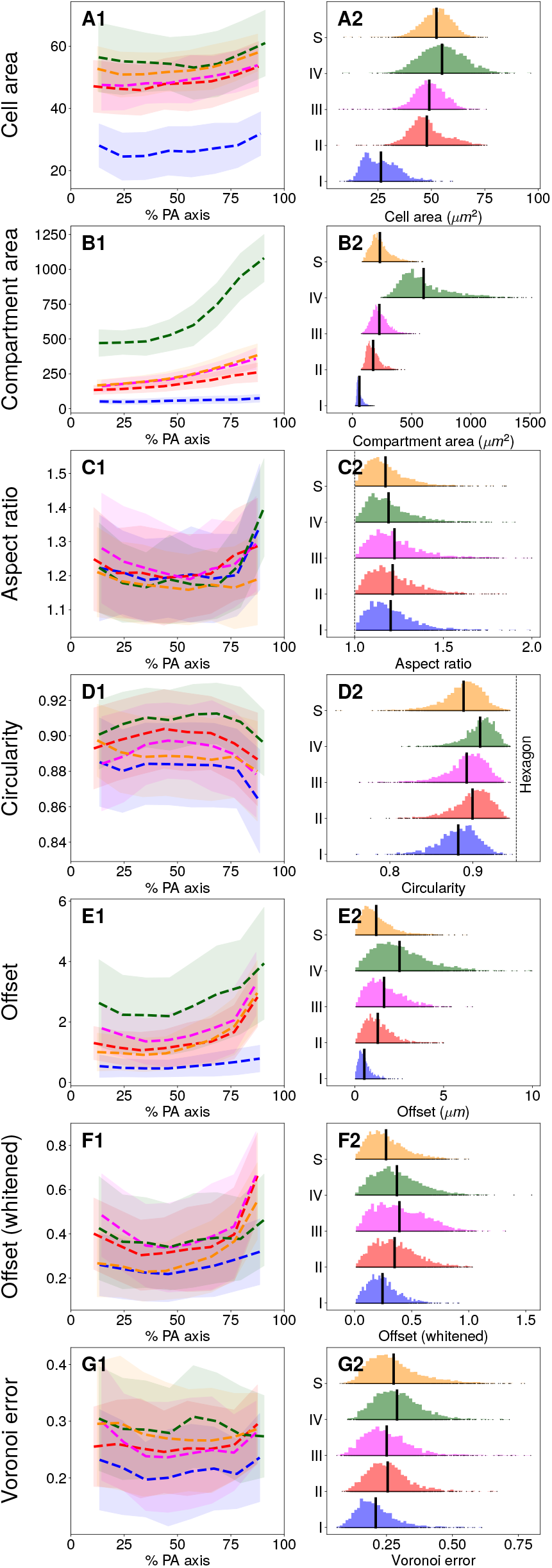
PA axis and life cycle variation. (A-G) Column 1: Computed metrics binned in 8 equally spaced segments along the PA axis. Means are shown as dashed lines with per-bin standard deviation reported by shaded segments. Colors correspond to developmental stages defined in Fig. 3. Column 2: Histograms of metrics by stage in 100 equally spaced bins, by stage, with empirical means indicated by vertical black bars. Units are noted in parentheses, and otherwise, are dimensionless.

In contrast, the CZ3 compartment area grows substantially along this axis, with a minimum increase of ∽50% in stage I and a maximum of ∽130% in stages III and Moreover, the slope increases after the equatorial region in all stages, an effect which is most pronounced in stage IV. Cell and compartment areas also increase by life cycle stage as shown in Fig. 8, rows A-B. Somatic cell areas double from I-II, growing merely ∽10% afterwards, whereas compartment areas expand primarily after III, with a ∽150% increase occurring from III-IV (SI Appendix, Table S2). This surge in compartment area mirrors the spheroid’s growth from III-IV, accounted an estimated ∽90% to parental ECM volume changes, ~ 10% to that of growing juveniles, and minimally to that of somatic cells (Tables 2 and SI Appendix, S1). Fig. 8A2-G2 shows distributions of the metrics; apart from cell area, all exhibit positive skew and exponential tails which suggest good fits with gamma-type distributions [39],

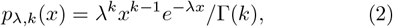

where *x* is suitably standardized. This skew should be considered when making mean-based comparisons across life cycle stages. The long left tails of cell area reflect the persistence of small somatic cells throughout the life cycle, confirmed by inspection in the chlorophyll signal. Lastly, the cell size distribution primarily translates rightward in time, while the compartment size distribution simultaneously translates and stretches, indicating an increase in polydispersity.

**TABLE 2.** Summary of estimated volumetric growth by stage. Estimated ECM volume is volume of spheroid minus that estimated of juveniles and somatic cells, as explained in SI Appendix, Table S1. Values in final two columns represent changes with respect to preceding stage.

| Stage | Parent radius ($\mu\text{m}$) | Offspring radius ($\mu\text{m}$) | Somatic cell radius ($\mu\text{m}$) | Parental ECM volume change (est., $\text{mm}^3$) | Parental ECM growth rate (est., $\text{mm}^3/\text{h}$) |
| --- | --- | --- | --- | --- | --- |
| I | $110 \pm 5.9$ | $16 \pm 2$ | $2.9 \pm 0.4$ | ↓ | ↓ |
| II | $220 \pm 19$ | $29 \pm 2$ | $3.9 \pm 0.3$ | 0.039 | 0.0026 |
| III | $240 \pm 13$ | $30 \pm 4$ | $3.9 \pm 0.3$ | 0.015 | 0.0025 |
| IV | $420 \pm 5.6$ | $79 \pm 6$ | $4.2 \pm 0.4$ | 0.23 | 0.014 |
| S | $260 \pm 26$ | $15 \pm 1$ | $4.1 \pm 0.3$ | n/a | n/a |

#### 2. CZ3 compartments transition from tighter polygonal to looser elliptical packing

While there is no apparent trend in the circularity of CZ3 compartments along the PA axis, the average circularity increases from stage I to IV. Since extracompartmental ECM space appears as compartments increase in circularity both effects correlate with enlargement of the spheroid. Fig. 8D2 shows that circularity increases in mean while decreasing in variance, suggestive of a relaxation process by which compartments of a particular aspect ratio but different polygonal initial configurations relax to a common elliptical shape with the same aspect ratio. This transition is also apparent by the ∽39% increase from stage I to IV in error with respect to the Voronoi tessellation (Fig. 8G2), whose partitions are always convex polygons.

#### 3. CZ3 compartments enlarge anisotropically

While the compartments become more circular as they expand, the aspect ratio is independent of stage and thus of organism size. The apparent increase at the ends of the posterior-anterior (PA) axis (U-shaped curves) is likely due to partially out-of-plane compartments appearing preferentially elongated. Fig. 8C shows that aspect ratio distributions are not only stable in mean, with less than 5% variation, but also in skewness and variance; they are gamma-distributed throughout growth (SI Appendix, Fig. S9). Together, the invariance of aspect ratio and increasing compartment circularity during growth suggests a transition from tightly packed, polygonal compartments (where neighboring boundaries are closely aligned) to elliptical configurations in which neighboring boundaries are no longer in full contact. We term this process *acircular relaxation*.

To study how ECM is distributed around the somatic cells, we quantified the cellular offset during the life cycle. The absolute offset from the compartment center of mass (Fig. 8E1-2) increases from stages I to IV, and along the PA axis, indicating a strong correlation between larger compartment areas and cellular displacements (as we show in § F). Perhaps counterintuitively, the cellular offset vector shows no correlation with the primary elongation axis of the compartment and is uniformly distributed in 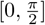 throughout I-IV and S (SI Appendix, Fig. S10). In contrast to the absolute offset, the whitened offset (which takes into account compartment size and anisotropy) is nearly constant in mean after stage II (black vertical lines in Fig. 8F). The support of the distribution does increase, albeit at a smaller rate than that of the cellular offset. Throughout this analysis of variation along the PA axis (Fig. 8), similarly sized spheroids in the asexual and sexual life cycle stages, bearing embryos or egg cells respectively, resemble each other in ECM geometry.

### F. CZ3 geometry shows feature correlations

The analysis above indicates compelling correlations between ECM growth and geometry. Here we analyze these in more detail with pooled data from all spheroids presented in Fig. 8. Figure 9A shows an exponential increase in the compartment area *a*_cz3_ with cell area *a*_cell_ through stage III, saturating at stage IV. This confirms that somatic cells primarily grow after hatching and before stage II, in contrast to compartments, which primarily grow after stage III.

**FIG. 9.**
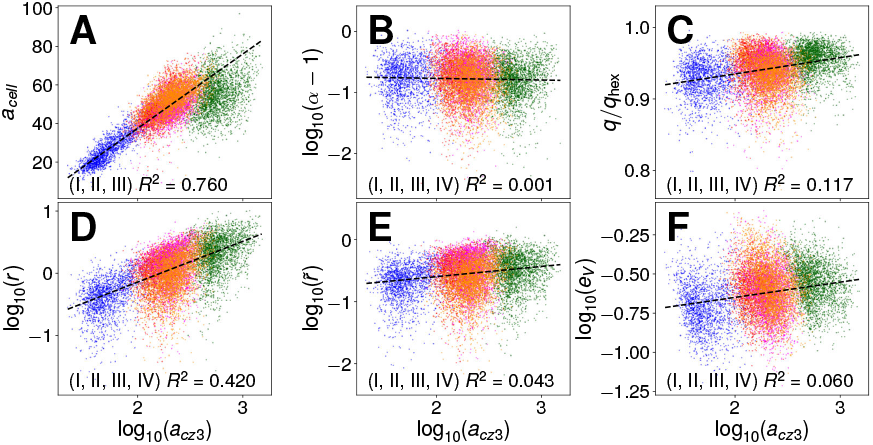
Pair correlations of compartment features. At stages I-IV (blue, red, magenta, green) and S (orange), plots show correlations between compartment area (*a*_cz3_) and metrics in Table 1 (*q*_hex_ in (C) is the circularity of a regular hexagon). Coordinate transforms are chosen in either linear-or log-scale, with natural offsets, to produce most equally-sized contours across the distributions as measured via the kernel density estimate. *R*^2^ is the linear regression correlation coefficient for stages listed.

As expected from the PA analysis, the aspect ratio (B) is decoupled from compartment size, while the circularity (C) increases. This reinforces the conclusion that as compartments expand they preserve their aspect ratio while decreasing in polygonality. The cellular offset (D) reveals a power-law relationship with compartment area, which, in conjunction with the weak coupling between whitened offset and compartment size (E), further supports this conclusion in light of a scaling argument presented in Discussion §5.

### G. Tessellation properties change during the life cycle

The metrics in Fig. 10 reveal clear trends by life cycle stage for the global geometry of each spheroid. Panel E shows the increasing circular radius, with most of the growth occurring between stages I-II and III-IV. II-III is separated by fewer hours and occurs during the first dark phase. Stage S is sorted in size close to stage III, supporting its resemblance with stages II and III in the PA analysis.

**FIG. 10.**
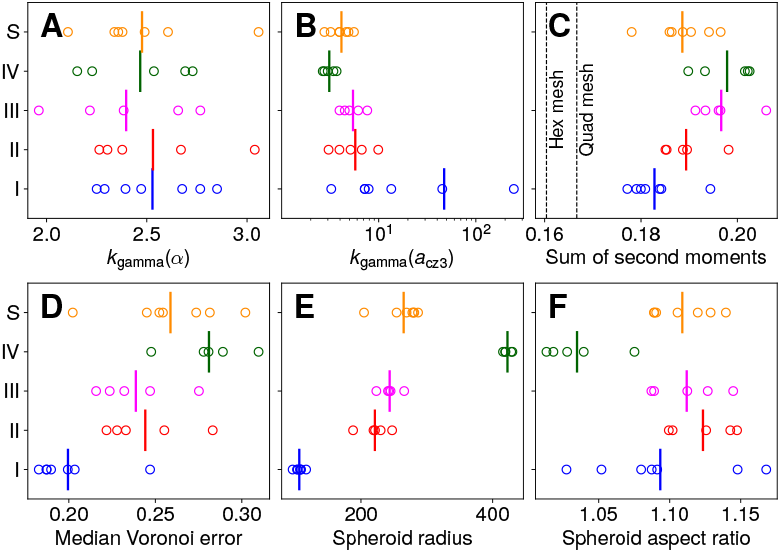
Global geometric properties of spheroids in stages I-IV, S. (A) shape parameter *k* of gamma distribution fit to *α*, (B) the same for *a*_*cz*3_, (C) sum of second moments (3), (D) median *e*_*V*_ over the whole spheroid, (E) the circular radius of each spheroid (geometric mean of major and minor axes), and (F) the aspect ratio of each spheroid (ratio of major and minor axes).

At fixed mean, the shape parameter *k*_gamma_ of the gamma distribution (2) is a measure of the entropy of the configuration, with high *k* indicating an increasingly crystalline, Gaussian-distributed configuration by the central limit theorem [42]. Fig. 10A confirms the stability of the aspect ratio distribution between stages, exhibiting values of *k*_gamma_ between ∽ 2™ 3, similar to ranges previously reported for confluent tissues [43]. Simultaneously, panel B shows that *k*_gamma_ in the distribution of *a*_cz3_ is decreasing in stages I-IV, so the configuration (primarily the anterior hemisphere, SI Appendix, Fig. S11) becomes increasingly disordered. The initial high values of *k*_gamma_ are consistent with the earlier observation that CZ3 compartments begin in a tightly-packed configuration, and as *k* quantifies regularity we infer that both tight packing and proximity to an equal-area lattice describes the initial configuration. The values of *k*_gamma_ between 2 and 3 in Stage IV are close to those for the Voronoi tessellations [39]. This has implications for regime of validity of the Voronoi approximation, as discussed in § E 2.

The standardized sum of second moments, defined as

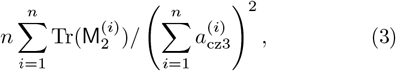

with *i* = 1 … *n* indexing CZ3 compartments per spheroid, is a space-partitioning cost that is minimized by equilateral hexagonal meshes (e.g. the surface of a honeycomb, see SI Appendix, §2.C.2). Although CZ3 compartments do not tile space due to extracompartmental spaces, (3) can nevertheless be computed for the covered area. This energetic cost for each spheroid, encoding deviations from optimality arising from some form of perimeter excess, is displayed in panel C. We find that somatic CZ3 configurations become *decreasingly* optimal during expansion—an observation consistent with the underlying increasing trends in cell offset and area polydispersity observed in § E 1 and § E 3. The metrics in B-C thus show the counterintuitive result that the space partitioning formed by CZ3 compartments becomes increasingly disordered as the spheroidal shape is maintained during the dramatic enlargement.

### H. Pherophorin II is also localized in the boundary zone

Confocal cross sections reveal that PhII is also part of the boundary zone (BZ), the outermost ECM layer of the organism (Figs. 1B and 4C and D). The PhII:YFP-stained BZ extends as a thin ∽ 1.1± 0.4 *µ*m arcing layer from the flagella exit points of one cell to those of all neighboring cells. This shape indicates that the outer surface of the spheroid has small indentations at the locations of somatic cells, where flagella penetrate the ECM. At these points, the BZ is connected to the CZ3 of the somatic cells below. Because the BZ is thin and not flat, it is not visible in a top, cross-sectional view of a spheroid through the centers of the somatic ECM compartments (e.g. Fig. 5), and only partly visible when the focal plane cuts through the BZ. If the focal plane is placed on the deepest point of the indentations, only the areas at which the BZ is connected to CZ3 can be seen (Fig. 11). From the centers of these areas the two flagella emerge and the flagellar tunnels are seen as two black dots due to the lack of fluorescence there (Fig. 11B). A closer look at the fine structure at the BZ-CZ3 connection site shows that fiber-like structures radiate from there to the BZ-CZ3 connection sites of neighboring somatic cells.

**FIG. 11.**
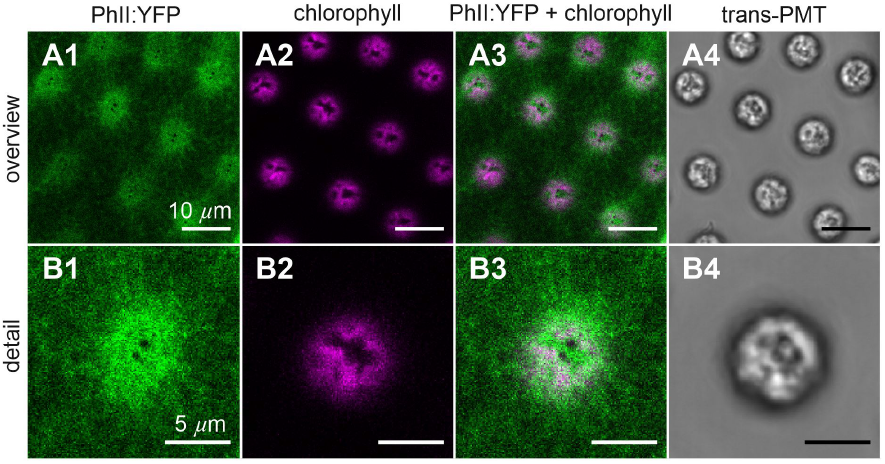
Magnified top view in regions where the BZ is connected to CZ3 in early stage II. Imaging as in Fig. 4. Fiber-like structures radiate from these regions. Flagellar tunnels are seen in the centers of those areas as two dark dots.

**FIG. 12.**
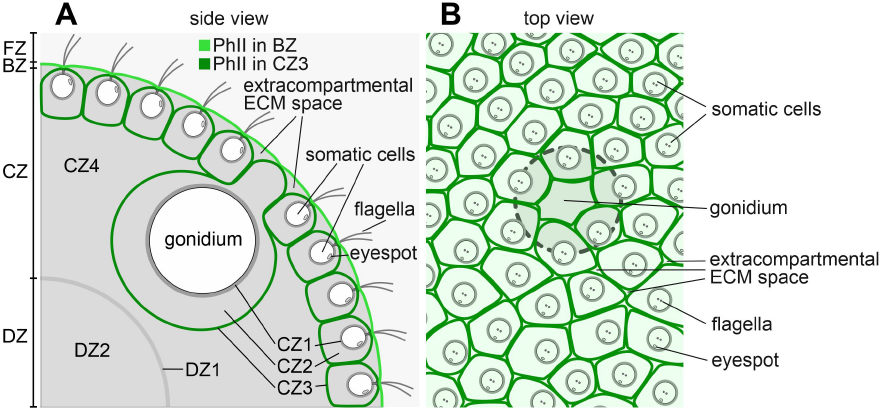
Overview of PhII:YFP localization in the ECM in early stage II. (A) Schematic cross section, showing localization in CZ3 of both somatic cells and gonidia (dark green) and BZ (light green). (B) Schematic top view, looking through the boundary zone, showing PhII:YFP in the CZ3 and the existence of extracompartmental ECM spaces. Position of a gonidium below somatic cell layer is indicated (dashed); the CZ3 compartment of the deeper gonidium extends to the surface where it is surrounded by ECM compartments with somatic cells.

## DISCUSSION AND CONCLUSIONS

### 1. Holistic view of PhII localization

Synthesizing the results of preceding sections, we arrive at the summary shown in Fig. 12 of the identified locations of PhII. It forms compartment boundaries (CZ3) not only around each somatic cell but additionally around each gonidium, and is also found in the outer border (BZ) of the spheroid; CZ3 and BZ are connected where the flagella emerge. While each compartment boundary can be assigned to the cell it encloses, and is most likely synthesized solely by that cell, PhII in the BZ is evidently formed collectively by neighboring cells. Since neither the compartment boundaries (CZ3) of the somatic cells are completely adjacent to the neighboring compartment boundaries, nor does the BZ rest directly on the compartment boundaries, extracompartmental ECM space remains between the CZ3 enclosures as well as between them and the BZ. The extracompartmental ECM space thus appears to be a net-like coherent space connected to CZ4.

### 2. Relation to earlier ECM studies by electron

#### microscopy

In earlier transmission electron microscopy images showing heavy metal-stained sections of the ECM, both the CZ3 and the BZ can be recognized as relatively dark structures, whereas the CZ2, CZ4 and the deep zone are very bright [20]. As the degree of darkness reflects the electron density and atomic mass variations in the sample, PhII evidently forms firmer wall-like structures in CZ3 and BZ, while CZ2, CZ4 and the deep zone have a very low density and are presumably of gel-like consistency. Using quick-freeze/deep-etch electron microscopy, it was shown that the fine structure of the ECM of volvocine algae such as *Chlamydomonas* and *Volvox* resembles a three-dimensional network [44, 45]. While both the CZ3 and BZ are likely dense networks with a fine pore size to the mesh, they must nevertheless allow the passage of small molecules and non-crosslinked ECM building blocks exported by cells, as evidenced by the growth during development of these compartments, the extracompartmental space, and the deep zone. Cells must also be able to absorb nutrients from the outside, which must pass through both the BZ and CZ3. The BZ may be a denser network than the CZ3, in order to prevent ECM building blocks from escaping into the environment.

### 3. Mechanical implications of identified ECM structures

As revealed by the localization of PhII:YFP and prior electron microscope studies, the BZ appears to form a dense “skin” on the outer surface of the spheroid to which the CZ3 compartments are firmly attached at the flagella exit points. This point-like attachment allows the compartments to expand in all directions during growth. As the CZ3 compartments are densely packed and attached to the BZ, they would be expected to provide rigidity as a kind of “exoskeleton” of the alga. A simple experiment shows this feature: if many *Volvox* are pressed together inside the suspending liquid medium and then released, they elastically repel each other. And while it is clear that the constant expansion of the compartments by incorporating further ECM components allows the ECM to enlarge considerably, the precise mechanism for transporting the ECM components to their destinations remains an open question.

### 4. Evolution of the volvocine ECM and convergence of a monolayer epithelium-like architecture

The ECM of *V. carteri* evolved from the (cellulose-free) cell wall of a *Chlamydomonas*-like, unicellular ancestor [7]. The cell wall of extant *Chlamydomonas* species consists of an outer “tripartite” layer with a highly regular, quasi-crystalline structure and an inner more amorphous layer. In few-celled volvocine genera with a low degree of developmental complexity, such as *Pandorina*, the tripartite layer is partially split, so that its outer leaflet is continuous across the surface of the organism, while its inner leaflet still surrounds each individual cell body [46]. In larger, more complex volvocine algae (*Eudorina* to *Volvox*), the entire tripartite layer is continuous over the surface of the organism. In the genus *Volvox*, the outer layer has developed even further and the tripartite layer has become part of the boundary layer, the BZ, while the inner layer has evolved species-specific (CZ3) compartments [20]. The architecture of these CZ3 compartments shows certain parallels with the epidermis of most plant leaves [47] and even epithelia in animals [38], all of which possess an (initially) closely packed, polygonal architecture. Looking at the plasma membranes in animal epithelia, cellulose-based cell walls in epidermal cells of land plants and cellulose-free CZ3 structures of somatic *Volvox* cells as touching compartment boundaries, their packing geometry can be described using the same physical concepts [48]. Interestingly, they also share the presence of an adjacent thin ECM layer, which represents a kind of boundary in all of them: the cuticle (secreted by plant epidermis cells), the basal lamina (secreted by animal epithelial cells) and the BZ (secreted by somatic *Volvox* cells). The shared geometrical solution likely represents an example of convergent evolution driven by the common pressure to evolve a monolayer epithelium-like architecture with protective and control functions.

### 5. Characteristics of somatic CZ3 geometry

Examination of the stochastic geometry of the space partition formed by CZ3 walls, along with somatic cell positions, reveals four key findings. First, somatic cell growth occurs mainly between stages I-II, whereas CZ3 compartments grow mostly during III-IV (Fig. 8), indicating that ECM production, rather than somatic cell enlargement is the primary driver of visible compartment growth (Table 2). Their surface areas are well-approximated by gamma distributions ((2)) with anterior/posterior hemispheres exhibiting different values of *k* (SI Appendix, Fig. S8). Such area distributions arise in granular and cellular materials [49, 50]; it is novel to observe them for intrinsic structures of an ECM, with non-stationarity in *k* revealing A/P differentiation.

Second, the aspect ratio distributions are remarkably invariant throughout the life cycle (Figs. 8 and 9), and well-approximated by gamma distributions with stable *k* (SI Appendix, Fig. S9). Maintaining a fixed *α≈* 1.2 requires that each compartment enlarges anisotropically, in strong contrast with the trend in aspect ratio of the overall spheroid, which is both lower and decreases from stages III-IV to less than 1.05 (Fig. 10F). This lack of local-global coupling suggests that compartment anisotropy could be set by geometric constraints [43] in the cellular configuration prior to stage I and may explain how the organism maintains (and increases) its sphericity despite the strong non-uniformity in size and shape of its compartments. Shape variability in the form of gamma-distributed aspect ratios arises in a large class of epithelial tissues and inert jammed matter [43] which exhibit deviations from optimality in the sense of (3). The epithelium-like architecture formed by the CZ3 robustly falls into this class.

Third, the somatic cell offset increases steadily with compartment area (Fig. 8D) while the whitened offset remains relatively constant (Fig. 8E). Both observations suggest a growth-induced deformation in which space is locally dilated (in a tangent plane containing the compartment) as ℝ^2^↦ *ρ* ℝ^2^, *ρ*≥ 1, which indeed increases offsets while preserving whitened offsets (SI Appendix, §B.1). Such transformations also naturally preserve the aspect ratio, consistent with earlier observations. The cellular offset angle with respect to the principal stretch axis, on the other hand, is a priori unconstrained by these observations, and in fact we find that it is uniformly distributed in [0, *π/*2] (SI Appendix, Fig. S10). This highlights a stochastic decoupling between cell positioning and compartment shape, much like that previously observed between compartment and spheroid principal stretches. Again, this may be established before stage I and subsequently scaled by compartment growth—analogous to two points diverging on the surface of an expanding bubble. Importantly, *global* dilations of space, likening *Volvox* itself to a bubble, cannot produce the increasing compartment area polydispersity we observe during the life cycle, while *local* dilations as above, likening the CZ3 ‘epithelium’ to a spherical raft of heterogeneously inflating bubbles, can. In a continuum limit, such local dilations may be represented by conformal maps [51].

Lastly, the CZ3 ‘bubbles’ transition from a space *partition* to a *packing* whose fraction decreases from near-unity while ECM-filled extracompartmental spaces allow the compartments to relax from polygonal to elliptical shapes. Such transitions are reminiscent of the hydration of foams [52], in which adding liquid (representing migration of ECM across the CZ3 walls, § C) alleviates contact constraints. Unlike classical foams, however, which relax rapidly to circular equilibrium shapes due to weak adhesion, inter-compartment adhesion is likely stronger due to crosslinking and the formation of insoluble networks. Such surface tension-adhesion trade-offs can result in acircular equilibria with decreasing packing fraction [53], unearthing one plausible explanation for the remarkable stability of aspect ratios as the compartments and extracompartmental spaces swell during growth (Fig. 5). Notably, in contrast to typical assumptions of epithelial models [53, 54], the CZ3 ‘raft’ is a self-assembled extracellular structure whose rheological properties including tension and adhesion are not under direct control by cells. This highlights more broadly the need to probe the rheology of the ECM—perhaps using microrheological techniques akin to those applied in cytoskeletal studies [55]—to understand how stochastic local interactions give way to robust global structures. Understanding the self-organization of crosslinking ECM components will shed light on the principles underlying the intricate geometry and stochasticity observed in multicellular extracellular matrices.

## MATERIALS AND METHODS

### Strains and culture conditions

Female wild-type strains of *Volvox carteri* f. *nagariensis* were Eve10 and HK10. Eve10 is a descendant of HK10 and the male 69-1b, which originate from Japan. The strains have been described previously [56–59]. Strain HK10 has been used as a donor for the genomic library. As a recipient strain for transformation experiments a non-revertible nitrate reductase-deficient (*nitA*^−^) descendant of Eve10, strain TNit-1013 [60], was used. As the recipient strain is unable to use nitrate as a nitrogen source, it was grown in standard *Volvox* medium [61] supplemented with 1 mM ammonium chloride (NH_4_Cl). Transformants with a complemented nitrate reductase gene were grown in standard *Volvox* medium without ammonium chloride. Cultures were grown at 28°C in a cycle of 8 h dark/16 h cool fluorescent white light [62] at an average of ∽100 *µ*mol photons m^−2^ s^−1^ photosynthetically active radiation in glass tubes with caps that allow for gas exchange or Fernbach flasks aerated with ∽50 cm^3^/min of sterile air.

### Vector construction

The genomic library of *V. carteri* strain HK10 in the replacement lambda phage vector *λ*EMBL3 [63] described by [27] has been used before to obtain a lambda phage, *λ*16*/*1, with a 22 kb genomic fragment containing three copies of the *phII* gene [28]. A subcloned 8.3 kb *Bam*HI-*Eco*RI fragment of this lambda phage contains the middle copy, the *phII* gene B, used here. The 8.3 kb fragment also includes the *phII* promoter region, 5’UTR and 3’UTR and is in the pUC18 vector. An artificial *Kpn*I side should be inserted directly upstream of the stop codon so that the cDNA of the *yfp* can be inserted there. This was done by cutting out a 0.5 kb subfragment from a unique *Mlu*I located 0.2 kb upstream of the stop codon to a unique *Cla*I located 0.3 kb downstream of the stop codon from the 8.3 kb fragment, inserting the artificial *Kpn*I with PCRs, and putting the *Mlu*I-*Cla*I subfragment back to the corresponding position. The primers 5’GTAACTAAC-GAATGTACGGC (upstream of *Mlu*I) and 5’*ATC-GAT* TCAGGTACCTGGCCCCGTGCGGTAGATG were used for the first PCR and the primers 5’<u>GGTACC</u>**TGA**TTGCCGTAAGAGCAGTCATG and 5’TCTAGCCTCGTAACTGTTCG (downstream of *Cla*I) for the second PCR (The *Kpn*I side is underlined, the stop codon is shown in bold). One primer contains a *Cla*I (italics) at its 5’end to facilitate cloning. PCR was also utilized to add *Kpn*I sides to both ends of the *yfp* cDNA. In addition, a 15 bp linker sequence, which codes for a flexible pentaglycine interpeptide bridge, should be inserted before the *yfp* cDNA. The *yfp* sequence was previously codon-adapted to *C. reinhardtii* [64] but also works well in *V. carteri* [11]. Since this *yfp* sequence was already provided with the linker sequence earlier [65], the primers 5’GGTACC*GGCGGAGGCGGTGGC* ATGAGC and 5’<u>GGTACC</u>CTTGTACAGCTCGTC and a corresponding template could be used (the *Kpn*I side is underlined, the 15 bp linker is shown in italics). The resulting 0.7 kb PCR fragment was digested with *Kpn*I and inserted into the artificially introduced *Kpn*I side of the above pUC18 vector with the 8.3 kb fragment. All PCRs were carried out as previously described [66–68] using a gradient PCR thermal cycler (Mastercycler Gradient; Eppendorf). The final vector pPHII-YFP (Fig. 2) was checked by sequencing.

### Nuclear transformation of *V. carteri* by particle bombardment

Stable nuclear transformation of *V.carteri* strain TNit-1013 was performed as described earlier [69] using a Biolistic PDS-1000/He (Bio-Rad) particle gun [70]. Gold microprojectiles (1.0 *µ*m dia., Bio-Rad, Hercules, CA, USA) were coated according to earlier protocols [66, 67]. Algae of the recipient strain where co-bombarded with the selection plasmid pVcNR15 [71], carrying the *V. carteri* nitrate reductase gene, and the non-selectable plasmid pPhII-YFP. Plasmid pVcNR15 is able to complement the nitrate reductase deficiency of the recipient strain and therefore allows for selection of transformants. For selection, the nitrogen source of the *Volvox* medium was switched from ammonium to nitrate and algae were then incubated under standard conditions in 9 cm. diameter petri dishes filled with ∽35 ml liquid medium. Untransformed algae of the recipient strain die under these conditions due to nitrogen starvation. After incubation for at least six days, the petri dishes were inspected for green and living transformants.

### Confocal laser scanning microscopy

For live cell imaging of transformed algae, cultures were grown under standard conditions and induced with 10 ×*µ*l medium of sexually induced algae in a 10 ml glass tube. An LSM780 confocal laser scanning microscope was used with a 63 ×LCI Plan-Neofluar objective and a 10 Plan-Apochromat (Carl Zeiss GmbH, Oberkochen, Germany). The pinhole diameter of the confocal was set to 1 Airy unit. Fluorescence of the PhII:YFP fusion protein was excited by an Ar^+^ laser at 514 nm and detected at 520-550 nm. The fluorescence of chlorophyll was detected at 650-700 nm. Fluorescence intensity was recorded in bidirectional scan mode for YFP and chlorophyll in two channels simultaneously. Transmission images were obtained in a third channel by using a transmission-photomultiplier tube detector (trans-PMT). Images were captured at 12 bits per pixel and analyzed using ZEN black 2.1 digital imaging software (ZEN 2011, Carl Zeiss GmbH). Image processing and analysis used Fiji (ImageJ 1.51w) [72]. To verify the signal as YFP fluorescence, the lambda scan function of ZEN was used in which a spectrum of the emitted light is generated by a gallium arsenide phosphide QUASAR photomultiplier detector that produces simultaneous 18-channel readouts. Emission spectra between 486 and 637 nm were recorded for each pixel with a spectral resolution of 9 nm using a 458/514 beam splitter and 488-nm laser light for excitation. After data acquisition, spectral analysis was performed to allow separation of heavily overlapping emission signals.

### Data, Materials, and Software Availability

All data and code are available on Zenodo (DOI: 10.5281/zenodo.14066435) [73].

## ACKNOWLEDGEMENTS

We are grateful to Kordula Puls and Diana Thomas-McEwen for technical assistance, and to Jane Chui and Kyriacos Leptos for inspiring conversations. Financial support for the work carried out in Bielefeld was provided by A.H.’s institutional funds. REG gratefully acknowledges the financial support of the John Templeton Foundation (#62220). This work was also supported in part by the Cambridge Trust (AS), and Wellcome Trust Investigator Grants 207510/Z/17/Z (SSMHH and REG) and 307079/Z/23/Z (SKB, SSMHH and REG).

## SI Appendix

### 1. SUPPLEMENTARY DATA: PHEROPHORIN II OVERVIEW AND DNA SEQUENCES

**FIG. S1.**
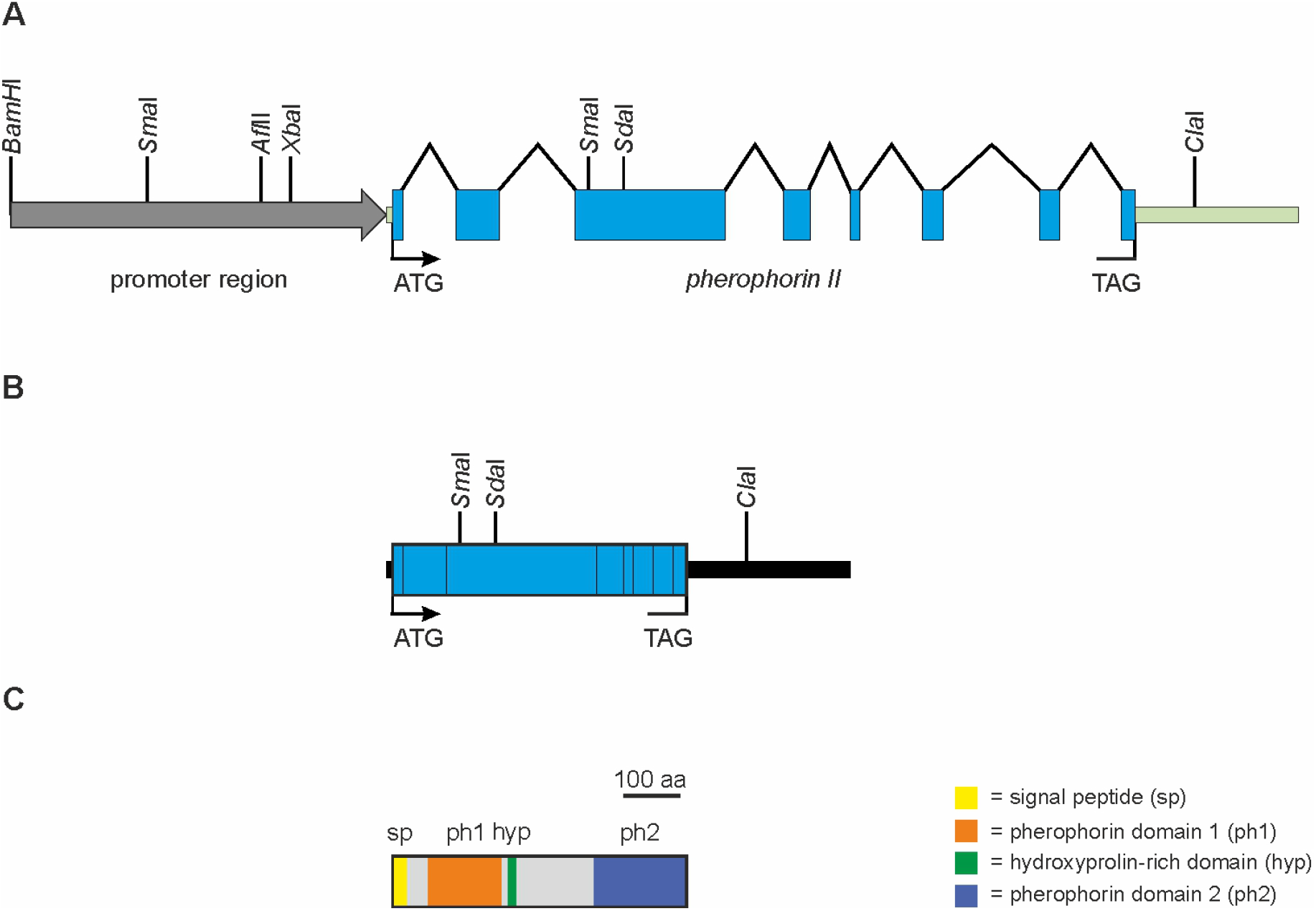
Schematic structure of the *phII* gene, *phII* mRNA and pherophorin-II protein. (A) The genomic region schematized here corresponds to the 8329-bp genomic fragment utilized in plasmid pPhII-YFP. The *phII* gene [S1] is located on scaffold 34 (nucleotides 980223 to 985045) of the *V. carteri* genome version 2.1 [S2] in Phytozome v13 [S3] on the reverse strand. The start codon is at nucleotide position 985025-985027 on the reverse strand. In the current *Volvox carteri* genome annotation available at Phytozome v13 (Volvox v2.1) pherophorin II is not annotated. Therefore the gene structure was established based on older annotations and confirmed with RNA-Sequencing data [S4]. The gene structure is indicated as follows: Coding sequences are represented by blue squares, intron sequences by carats, UTRs by green bars and the promoter region by a grey arrow. Start (ATG) and stop (TAG) codons are highlighted. The given restriction sites are also marked in SI Appendix, Fig. S2, which represents the genomic sequence of the *phII* gene. (B) Structure of the *V. carteri phII* mRNA. Sequence features are as indicated in A. The coding sequence (blue squares) totals 1557 nucleotides. The 5’ UTR is 18 bp in length, while there is a quite long 3’ UTR of 869 bp. The complete mRNA is 2,444 nucleotides in length. (C) Structure of the *V. carteri* pherophorin-II protein. The polypeptide comprises 518 amino acids and the calculated molecular weight amounts to 54.5 kDa. As pherophorin-II is an extracellular protein, it possesses a cleaved N-terminal signal peptide (sp) of 24 amino acids. In the mature protein three domains can be identified: an N-terminal pherophorin domain (ph1) with an E-value of 4.5e-27, a hydroxyproline-rich domain (hyp) in the middle [S5] and a C-terminal pherophorin domain (ph2) with an E-value of 2.1e-32.The pherophorin domains where identified by blasting the Pfam database [S6] using the hmmscan function [S7]. The short Hydroxyproline-rich domain consists of 78% (hydroxy) prolines (seven of nine amino acids).

**FIG. S2.**
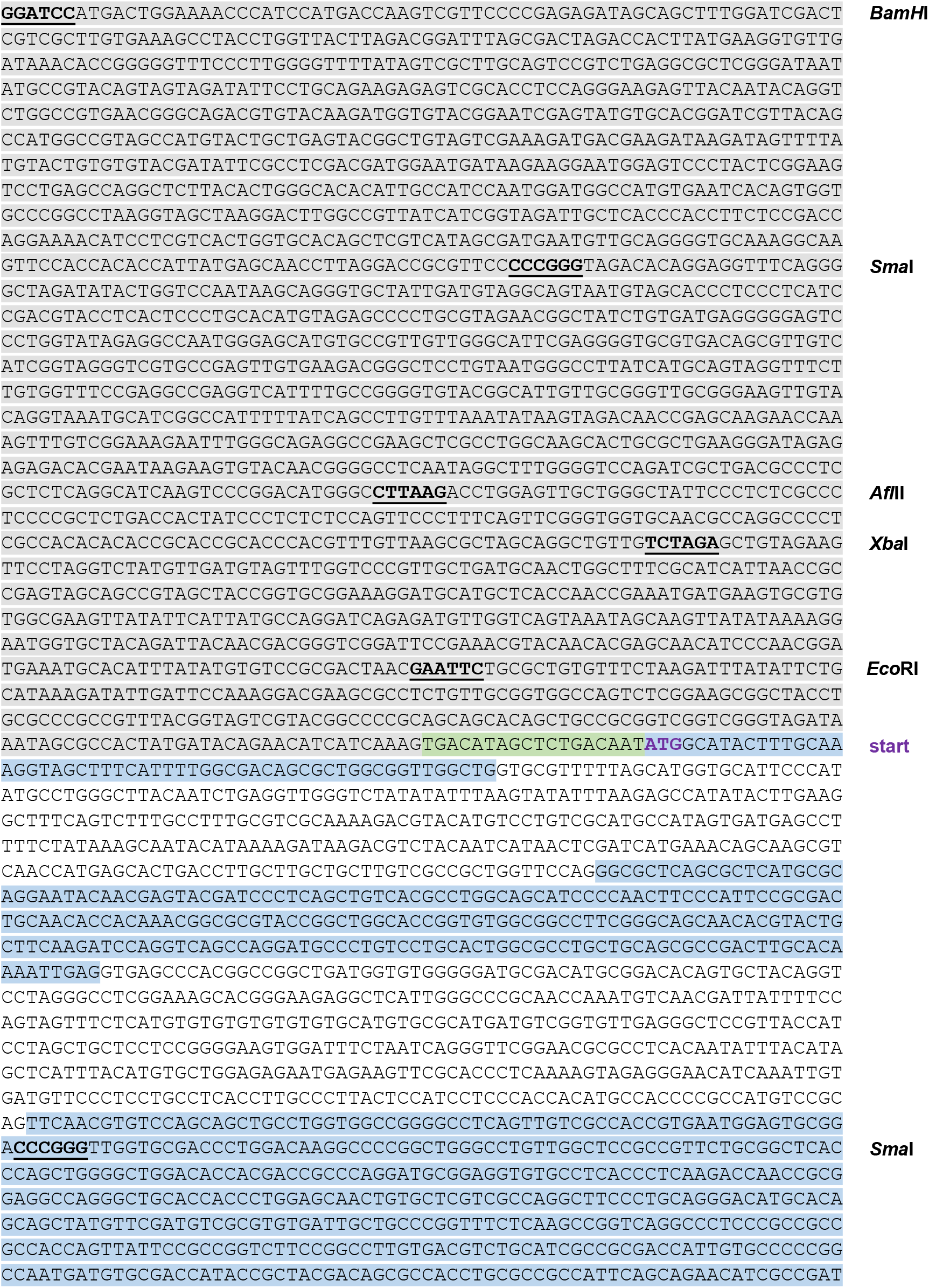
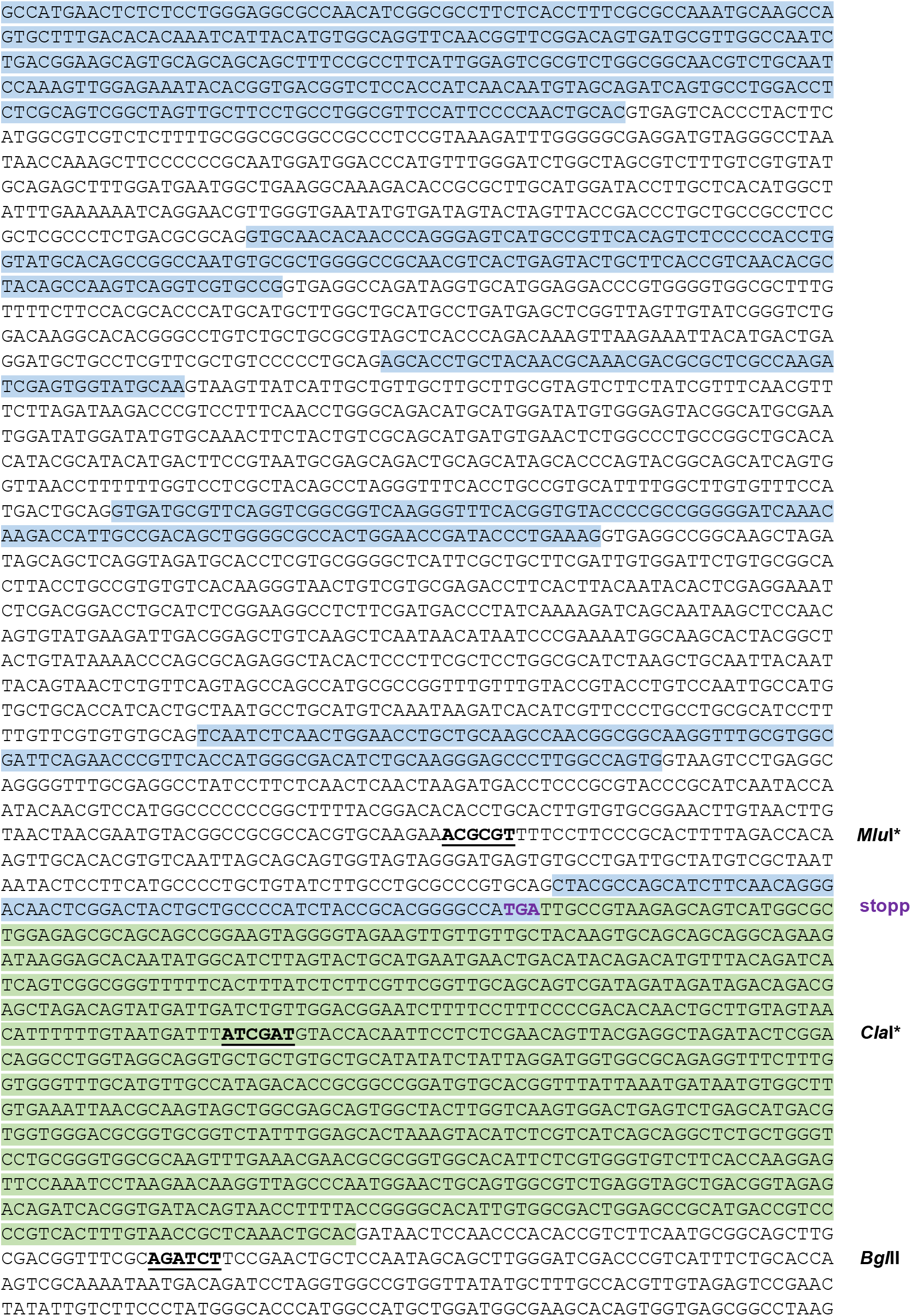
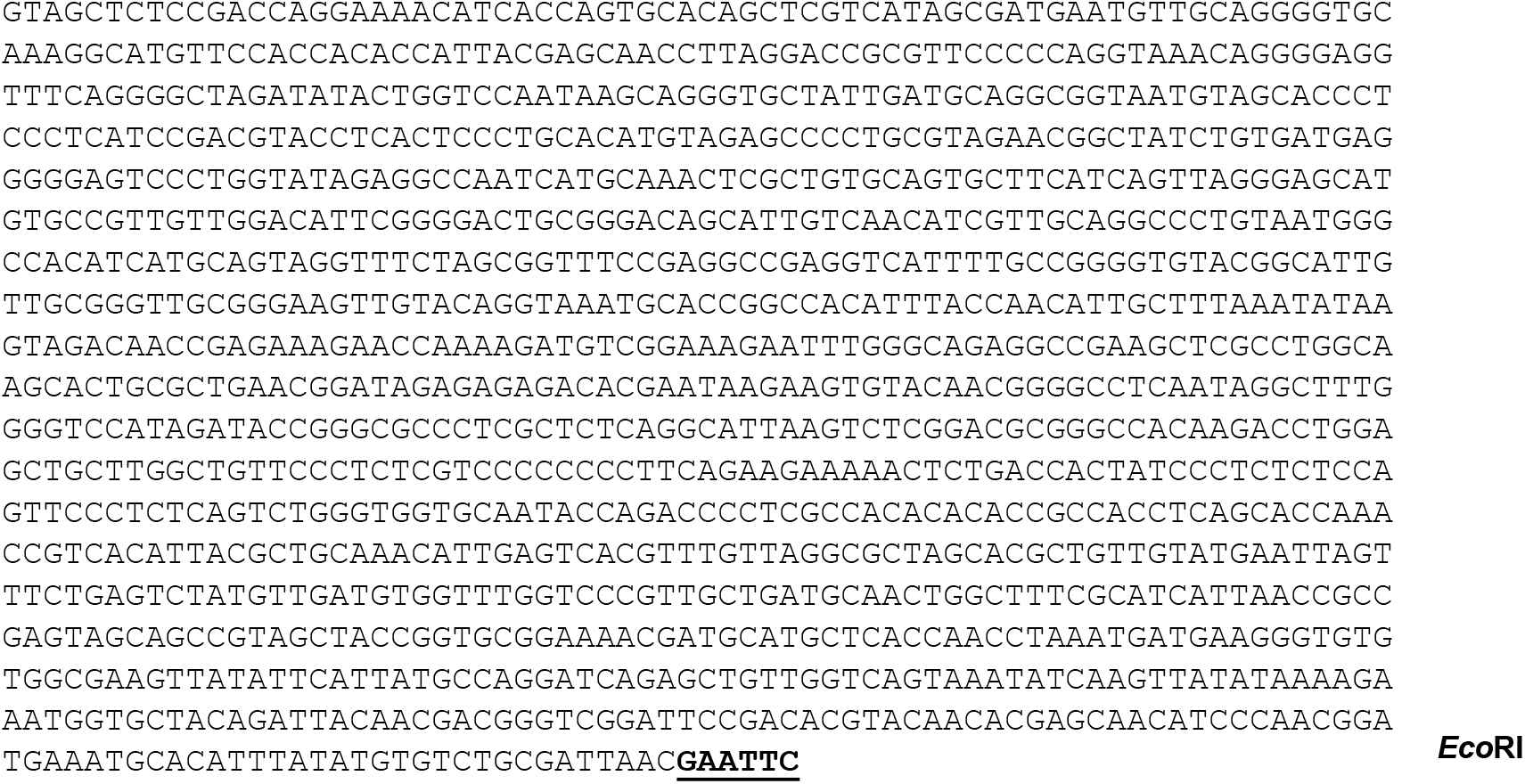
Genomic sequence of the *V. carteri phII* gene. The sequence depicted here corresponds to the 8329-bp genomic fragment utilized in plasmid pPhII-YFP. The *phII* gene [S1] is located on scaffold 34 (nucleotides 980223 to 985045) of the *V. carteri* genome version 2.1 [S2] in Phytozome v13 [S3] on the reverse strand. The start codon is at nucleotide position 985025-985027 on the reverse strand. In the current *Volvox carteri* genome annotation available at Phytozome v13 (Volvox v2.1) pherophorin II is not annotated. Therefore, the gene structure was established based on older annotations and confirmed with RNA-Sequencing data [S4]. The gene structure is indicated as follows: Coding sequences are shown with blue background, UTRs with green background and the promoter region with grey background. Start and stop codons are highlighted (violet font). The 5’ UTR is just 18 bp in length, while there is a quite long 3’ UTR of 869 bp. The coding sequence totals 1557 nucleotides. The restriction sites that are shown in Figure 2 (main text) and SI Appendix, Fig. S1 A and B are marked (bold, underlined). Restriction sides that were used for inserting the *yfp* coding sequence are marked with asterisks.

**FIG. S3.**
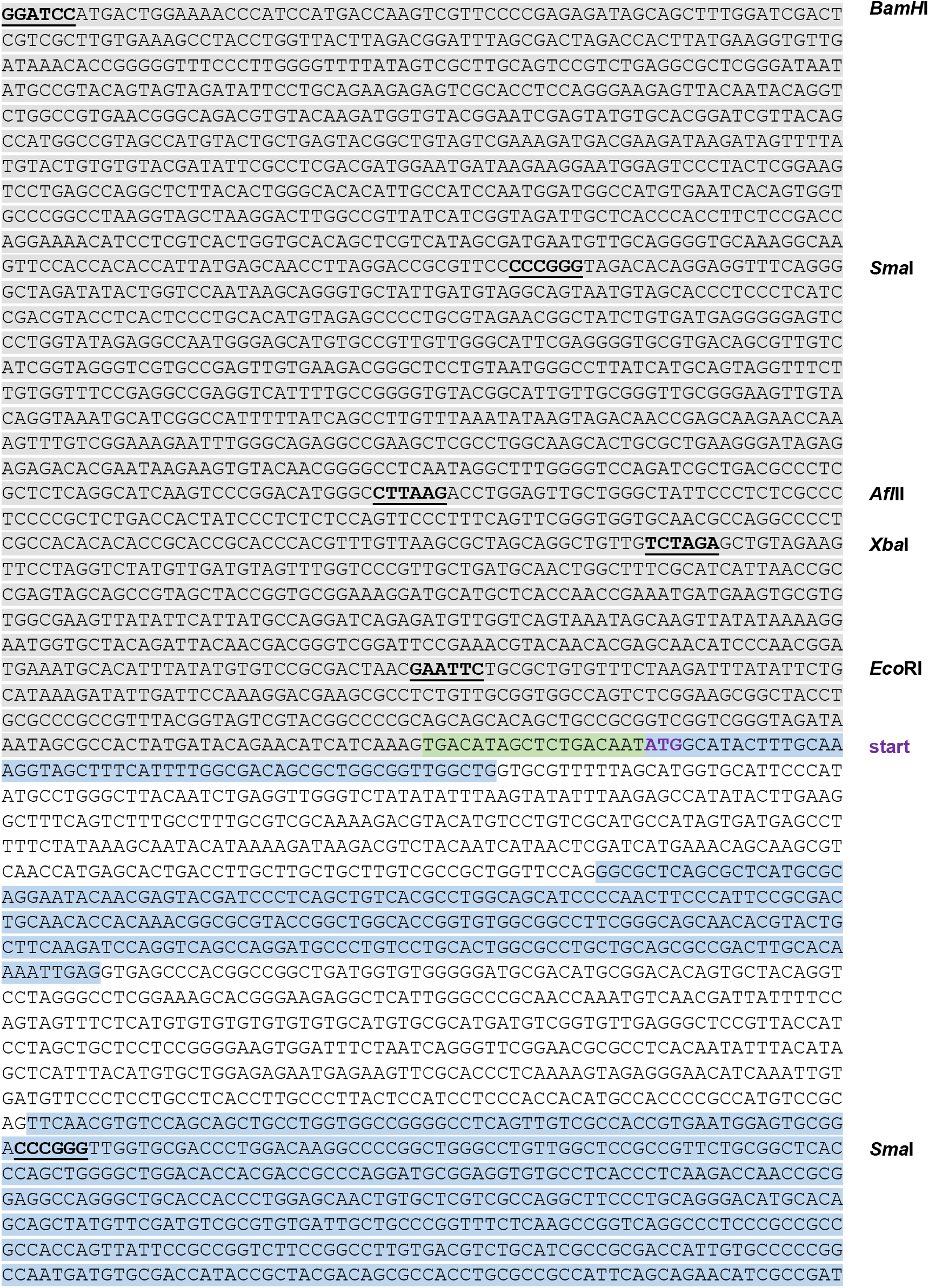
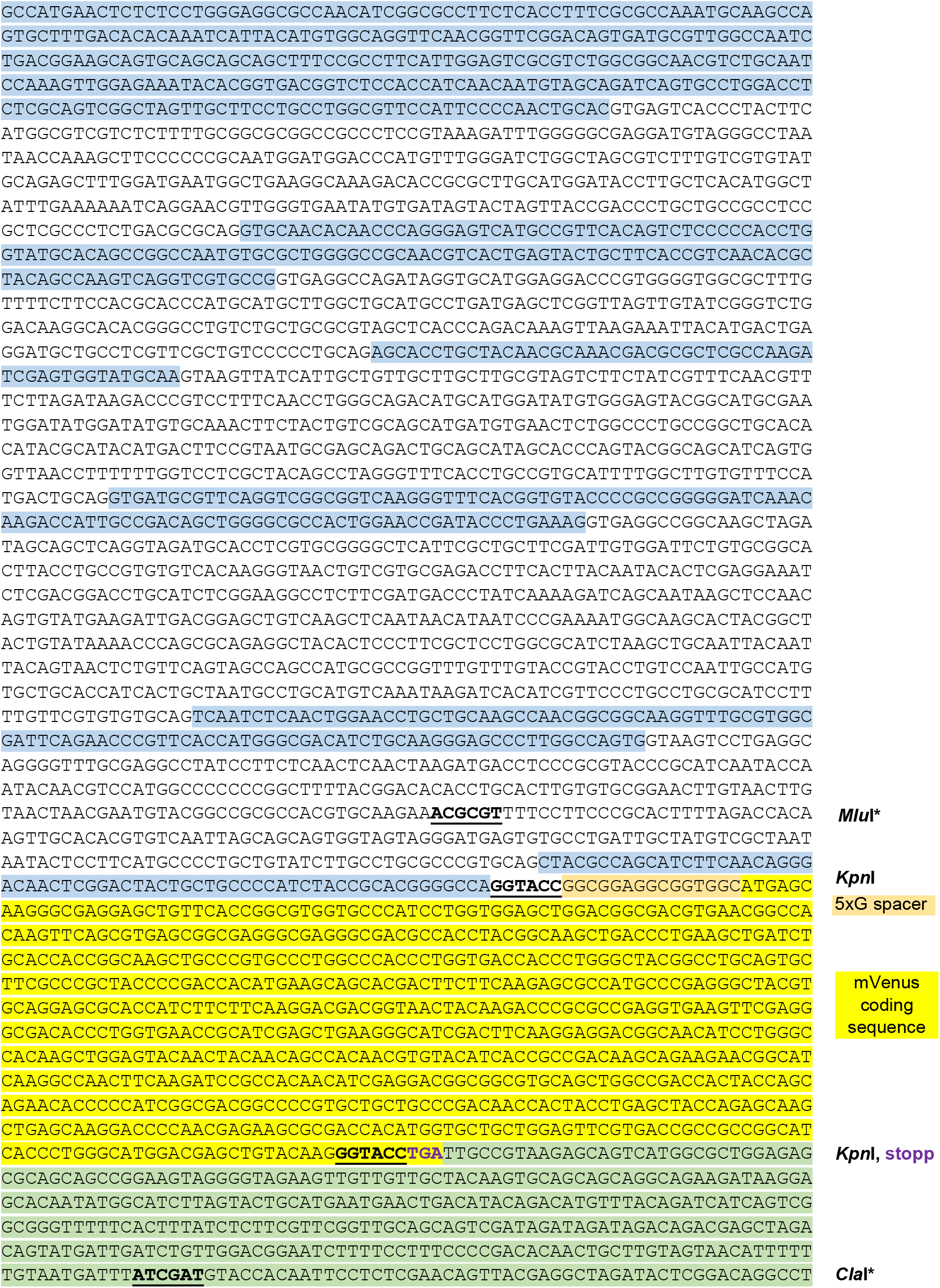
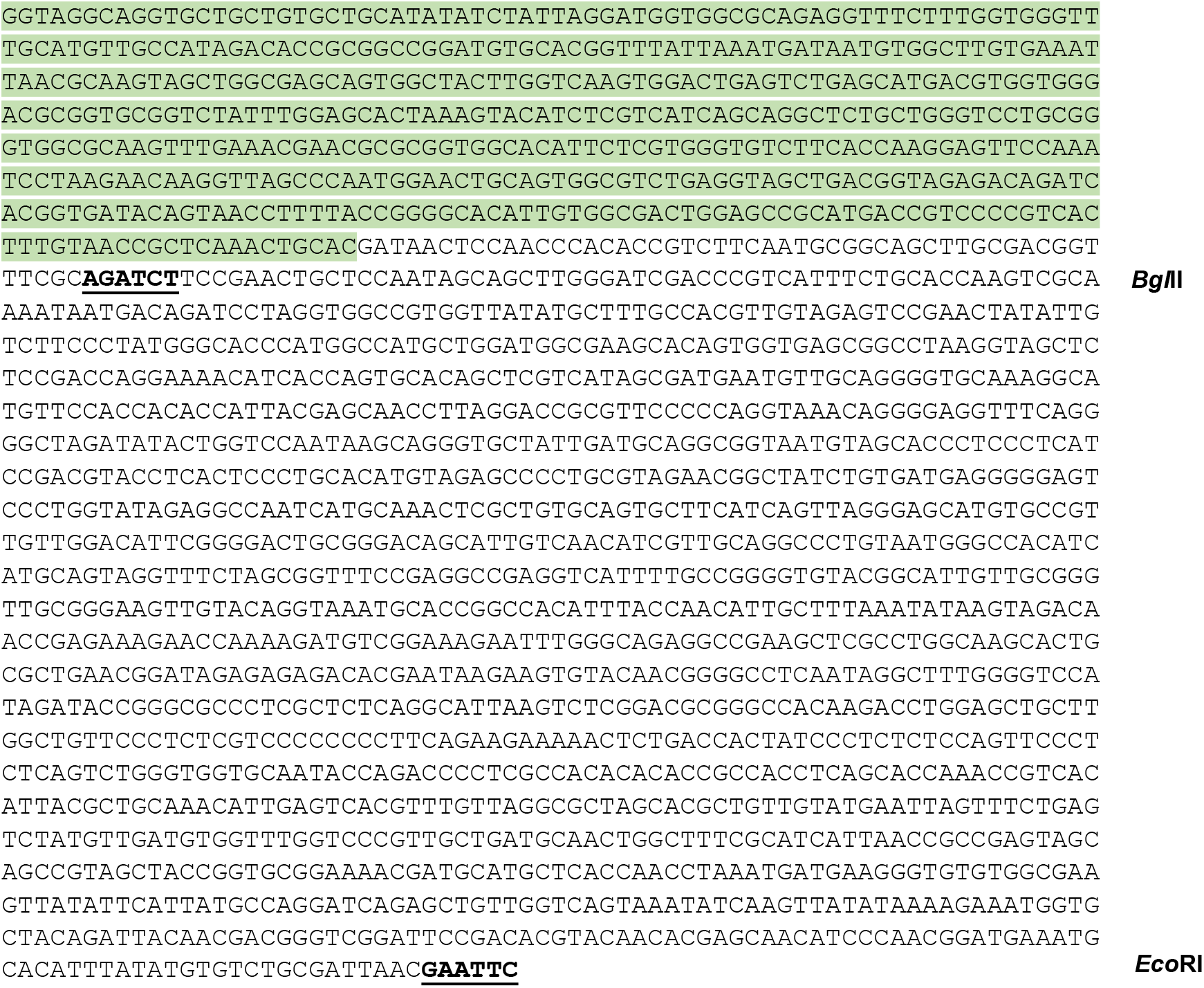
Genomic sequence of the *V. carteri phII* gene with *yfp* coding sequence added. The sequence depicted here is present in plasmid pPhII-YFP. The chimeric gene consists of 8.3 kb genomic DNA including 5’ and 3’ regulatory sequences and the complete transcribed region of pherophorin II including all introns as present on scaffold 34. The *yfp* (mVenus) [S8] coding sequence was fused in frame with the coding sequence of the last *phII* exon and a 15 bp sequence coding for a pentaglycine flexible spacer. For the insertion of mVenus, artificial *Kpn*I sides were introduced by recombinant PCR which also allow for a later exchange of the fluorescent marker. The gene structure is indicated as follows: Coding sequences are shown with blue background, UTRs with green background and the promoter region with grey background. Start and stop codons are highlighted (violet font). The 5’ UTR is just 14 bp in length, while there is a quite long 3’ UTR of 1,168 bp. The coding sequence of mVenus is highlighted in yellow. The pentaglycine spacer is highlighted in orange. The restriction sites that are shown in SI Appendix, Fig. S1 A and B are marked (bold, underlined). Restriction sides that were used for inserting the *yfp* coding sequence are marked with asterisks.

### 2. SUPPLEMENTARY METHODS: SEMI-AUTOMATED IMAGE SEGMENTATION AND GEOMETRIC ANALYSIS

#### A. Overview

We employ a semi-automated image analysis pipeline which uses Cellpose [S9] as a key step.

1. Contrast stretching of the image is performed by predetermined cutoffs, e.g. 2nd and 98th percentile intensity.
2. A J-invariant filtration is performed using (depending on the channel, fluorescence or trans-PMT) either (i) total-variation (TV) denoising by minimizing the Rudin-Osher-Fatemi functional

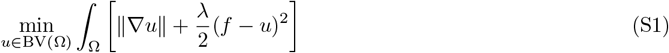

where *f* is the intensity profile of an image supported on the image domain Ω to be denoised, or (ii) wavelet denoising with adaptive thresholding. A J-invariant filter is defined as one whose output value at every pixel is independent of the value of the source pixel (i.e. is only a function of source pixels at other locations). We find TV denoising in particular to be effective at filtering Poisson noise and significantly enhance the performance of Cellpose on low-SNR fluorescence images. We did not use the trained denoising model available in Cellpose3 [S10].
3. A user-prompted input polygon *P* is used to estimate the diameter *d* = {∥max ***v***_*i*_ ***v***_*j*_∥ |***v***_*i*_, ***v***_*j*_∈ *P}* of typical instances to be identified in the image, passed as the *diameter* input to the Cellpose *cyto3* model [S10].
4. Objects identified as pixel-space masks by Cellpose are converted to polygons in the plane by either (i) taking the convex hull, for convex objects such as somatic cells, or (ii) identifying outlines in the mask. Degenerate and invalid polygons are suppressed by taking a single binary erosion-dilation step. Further conversion to ellipses for approximately elliptical objects (such as parent and offspring spheroids) is performed by computing the ellipse with the same *n*th-order moments of area as the polygon (see §2 B) up to *n* = 2.
5. False-positives are manually rejected where identified. False-negatives, where identified, are re-prompted to Cellpose by restricting to a user-specified region of interest around the object, and re-iterating from step 1.
6. For identification of the somatic CZ3 geometry in particular, only the somatic cell-CZ3 compartment pairs (as seen in Fig. 5, main text) which have jointly been successfully identified are retained.
7. Downstream analysis of the resulting polygons and/or ellipses is performed as described in Table 1 (main text) and further detailed below in §2 B.

#### B. Geometric moments of area

The geometric moments of bounded planar domains, analogous to the moments of bivariate uniform random variables, quantify shape properties such as size, center of mass, eccentricity, skew, and so on. In the case of polygonal domains, the geometric moments up to sufficiently high order completely determine the vertices [S11].

**Definition 2.1** (Planar *n*th area moment tensor). Let *D*⊂ ℝ^2^ be an open measurable set with boundary given by a simple closed curve ∂*D* = *C*. The *n*th moment tensor for the domain *D* of uniform mass density is

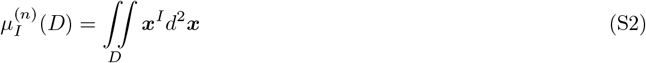

where *I* = (*i*_1_, …, *i*_*n*_) is a multi-index such that *i*_*j*_ ∈ {1, 2} and

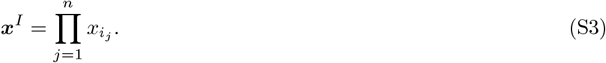

The 0th moment *µ*^(0)^ is the area of *D*. Accordingly, one may define the radius *R*_*D*_ of an equivalent (same-area) circle as 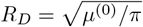.The *n*th-order moments can be calculated for polygons by (i) applying the divergence theorem to write (S2) as a boundary integral on *C*, and (ii) computing the integral as a finite sum using the piecewise-linearity of the sides *C*.

#### 1. First moment and centrality

**Definition 2.2** (Centroid). The center of mass of *D* is

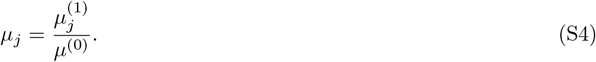

The notation ***µ*** = [*µ*_1_, *µ*_2_] evokes the probabilistic interpretation as the expected value of a uniform distribution supported on *D*. The basis in which (S2) is computed will unless otherwise specified be taken, for moments of order ≥ 2, to be one in which ***µ*** is at the origin.

**Definition 2.3** (Dimensionless centrality of a test point). For a test point ***y*** ∈ ℝ^2^, we define the centrality metric

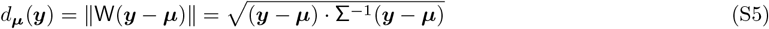

with W a matrix defined as W^⊺^W = ∑^−1^, and ∑ the covariance matrix of *D*, defined in (S7).

This *whitening* procedure (again evoking the probabilistic interpretation of ∑) enables comparison across domains *D* of varying second moment, resulting in a quantity which is dimensionless. In probability terms, (S5) is the Mahalanobis distance of ***y*** to the uniform distribution supported on *D*.

In particular, *d* is scale-invariant; for dilations of space ℝ^2^ → *ρ*ℝ^2^, *ρ >* 0, we have ***µ*** → *ρ****µ, y*** → *ρ****y***, and ∑ → *ρ*^2^∑ by (S7), hence *d*_***µ***_(***y***) → *d*_***µ***_(***y***) by (S5). Moreover, dilations preserve aspect ratios, since

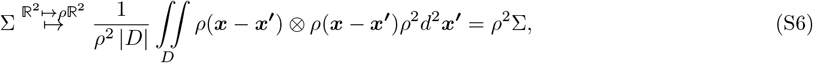

hence its eigenvalues map as *λ*_*j*_ → *ρ*^2^*λ*_*j*_ and the ratio *λ*_max_*/λ*_min_ is preserved.

#### 2. Second moment and isotropy

**Definition 2.4** (Covariance matrix of a domain). The normalized second central moment, or covariance matrix, is

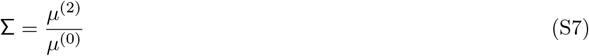

As before, *central* indicates that *µ*^(2)^ is computed in a basis in which ***µ*** is at the origin.

In probability terms, ∑ is the covariance matrix of a uniform distribution supported on the domain *D*. If *D* has radial symmetry about ***µ*** (either continuous, as a circle does, or discrete, as regular *n*-gons do), then ∑ = *c*I for some *c >* 0. This is verified by the existence of an eigenspace of dimension 2. ∑^−1^ = M is the matrix defining an ellipse with the same aspect ratio as *D*.

**Definition 2.5** (Principal axes of a domain). Using (S7), we may define the principal axes and stretches of a domain *D* via Hermitian eigendecomposition

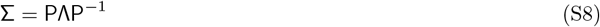

with ***v***_*i*_ the columns of P being the principal axes and and *λ*_*i*_ *>* 0 in ascending order (as ∑ is symmetric positive-definite for non-degenerate curves).

**Definition 2.6** (Aspect ratio of a domain). Let *λ*_1_, *λ*_2_ and *v*_1_, *v*_2_ be the principal stretches and axes (in ascending order) of ∑ as in (S8). The aspect ratio is

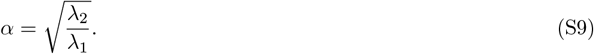

Ellipses (objects defined uniquely by their moments to order *n* = 2) with the same orientation, aspect ratio, and area (*πab*) as *D* have the minor and major axes

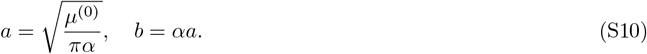

Equation (S10) is an exact elliptical representation of a polygon (or arbitrary domain), in contrast to elliptical approximations, e.g. (i) weighted *ℓ*^2^-minimization of vertex distance or (ii) convex programs for minimum-area bounding ellipses.

#### 3. Moments under affine transforms

Let F *>* 0 be a symmetric positive-definite matrix (e.g. a strain tensor) and define an affine transform of a domain *D* about its center of mass by

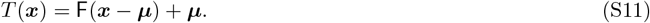

Let *T* (*D*) be the transformed region. By change of coordinates for integrals, the relevant moments of *T* (*D*) are

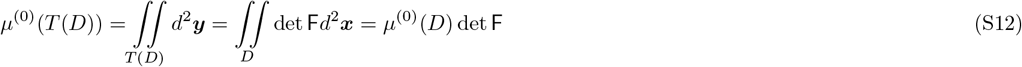

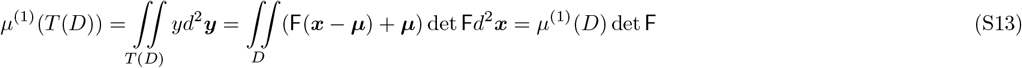

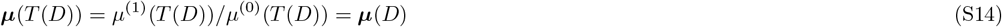

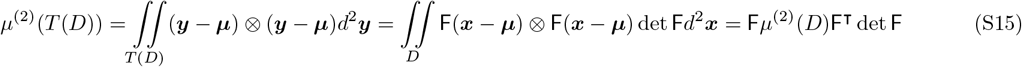

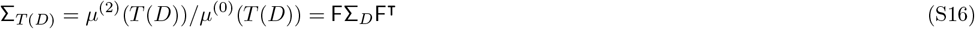

hence the centroid is preserved and the covariance matrix maps as ∑ → F∑F^⊺^.

#### 4. Whitening a domain

Let a domain *D* have covariance matrix ∑ (S7). Define the *whitened* domain *D*_*W*_ by the affine transform *T* as defined in (S11),

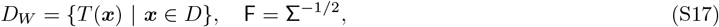

with the matrix square root F typically approximated by singular value decomposition (SVD) as

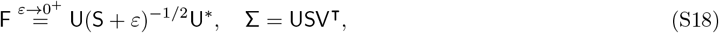

and F the *ZCA whitening matrix* [S12] with regularization constant *ε≪* 1. Then *D*_*W*_ has aspect ratio 1 as defined in (S9).

### C. Isoperimetric problems

We recall here several quantities which can be used as measures of the deviation of a domain *D* from a disk.

#### 1. Classical isoperimetric inequalities

One has

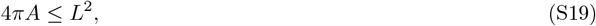

where *A* is the area of *D* and *L* the total arclength of *C* (which we may now require to be a rectifiable Jordan curve) and is an equality only for circles. Accordingly, one defines an *isoperimetric quotient*, which we term the *circularity* in the main text,

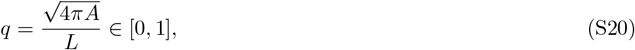

maximized for disks. For regular *n*-gons, one has

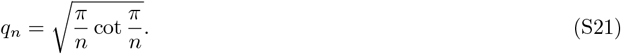

A natural question is whether (S19) can be used to define a set distance, in the sense of Hausdorff metric, of *D* to the “best” disk. This turns out [S13] to be related to the problem of making (S19) *quantitative*, in the sense of a nonnegative quantity ν(*D*) such that

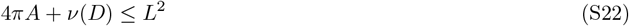

for all *D*, with ν(*D*) = 0 iff *D* is a disk. Without proof, we cite [S13] the result that the *isoperimetric deficit*, defined as the dimensionless quantity

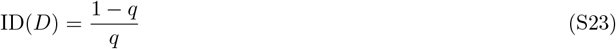

upper-bounds any such quantitative inequality ν(*D*) via

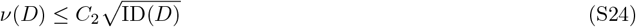

for some dimension-dependent constant *C*_2_. For convex *D* (applicable e.g. to the cells of a Voronoi tessellation), (S23) upper-bounds the Hausdorff distance *d*_*H*_ to the best-fit equal-volume ball *B* as

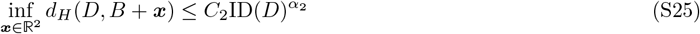

for dimension-dependent constants *C*_2_, *α*_2_ [S13]. Equality is achieved for *D* which is a ball.

#### 2. Weighted isoperimetric inequalities

Recalling the second moment *M*_2_ as defined in the main text, eq. 1, written here with explicit arguments *M*_2_(***x***, *D*), we recall by classical results that its minimization is also an isoperimetric problem.

**Lemma 1** (Disks minimize Tr(*M*_2_)). Let *D* ⊂ ℝ^2^ be as in Definition 2.1, |*D*| its Lebesgue measure, and ***x*** ∈ ℝ^2^. We have 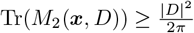 with equality iff *D* is a disk and ***x*** is its centroid.

*Proof*. Write Tr(*M*_2_) as

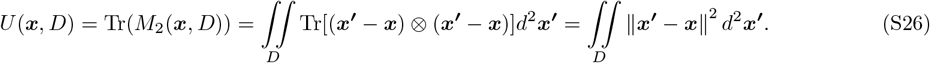

Then optimality in the first argument implies

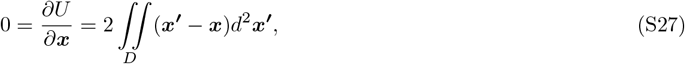

which holds iff ***x*** = ***µ*** is the centroid of *D*. Locating ***µ*** at the origin without loss of generality, we have

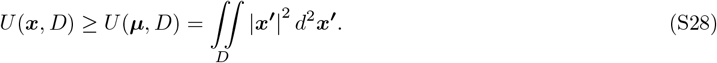

Recognizing the last expression as the polar moment of inertia and applying a *weighted* isoperimetric inequality (7.2, [S14]),

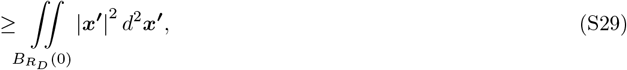

where 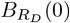 is a disk at the origin of the same area as *D* (i.e. *R*_*D*_ is *D*’s circular radius as in §B). Thus

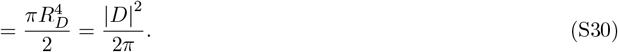

Recall also the (standardized) sum of second moments, Eq. 3, main text. Using the previous lemma we may establish the following optimality for it in terms of honeycombs.

**Lemma 2** (Second moment bound for tessellations). For any space packing 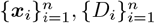 with standardized sum of second moments *m*_2_ defined as

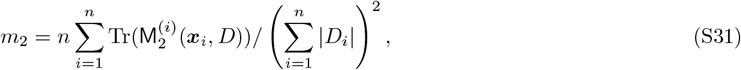

we have 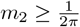.

*Proof*. 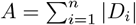and 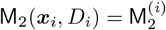 Then

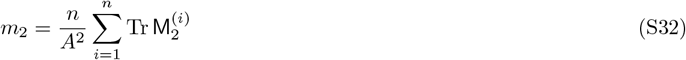

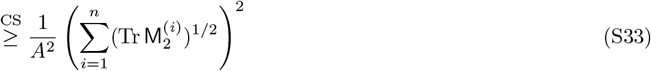

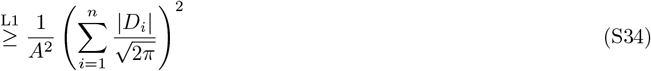

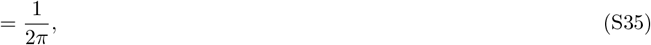

where the first inequality is Cauchy-Schwarz and the second is Lemma 1.

Note that the first inequality is sharp only for configurations consisting of congruent cells *D*_*i*_. The second inequality cannot be sharp for space *partitions*, only *packings* consisting of congruent disks. Space *partitions* optimize the second only if they are asymptotically (in *n*) regular hexagonal lattices [S15, S16].

#### D. Average number of neighbors in a spherical Voronoi tessellation

Let 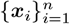 be a collection of points on the surface of a sphere and 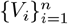 the corresponding set of spherical polygons given by their Voronoi tessellation. The number of neighbors of each *V*_*i*_ is precisely the degree (number of incident edges) *d*(***x***_*i*_) of each node in the topological dual, the Delaunay triangulation 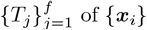.Let *m* be the number of edges; since this is a triangulation of a compact (boundaryless) surface, we have the relation 3*f* = 2*m*. By the Euler theorem,

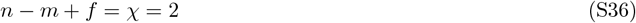

where *χ* is the Euler characteristic of the sphere. Substituting the triangulation property into (S36) yields *m* = 3*n™* 6, hence the average degree 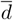 is

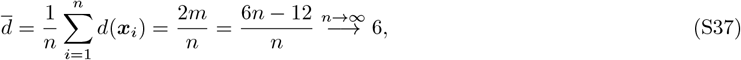

thus the average number of neighbors of a Voronoi polygon is asymptotically 6.

### 3. SUPPLEMENTARY ANALYSES

**FIG. S4.**
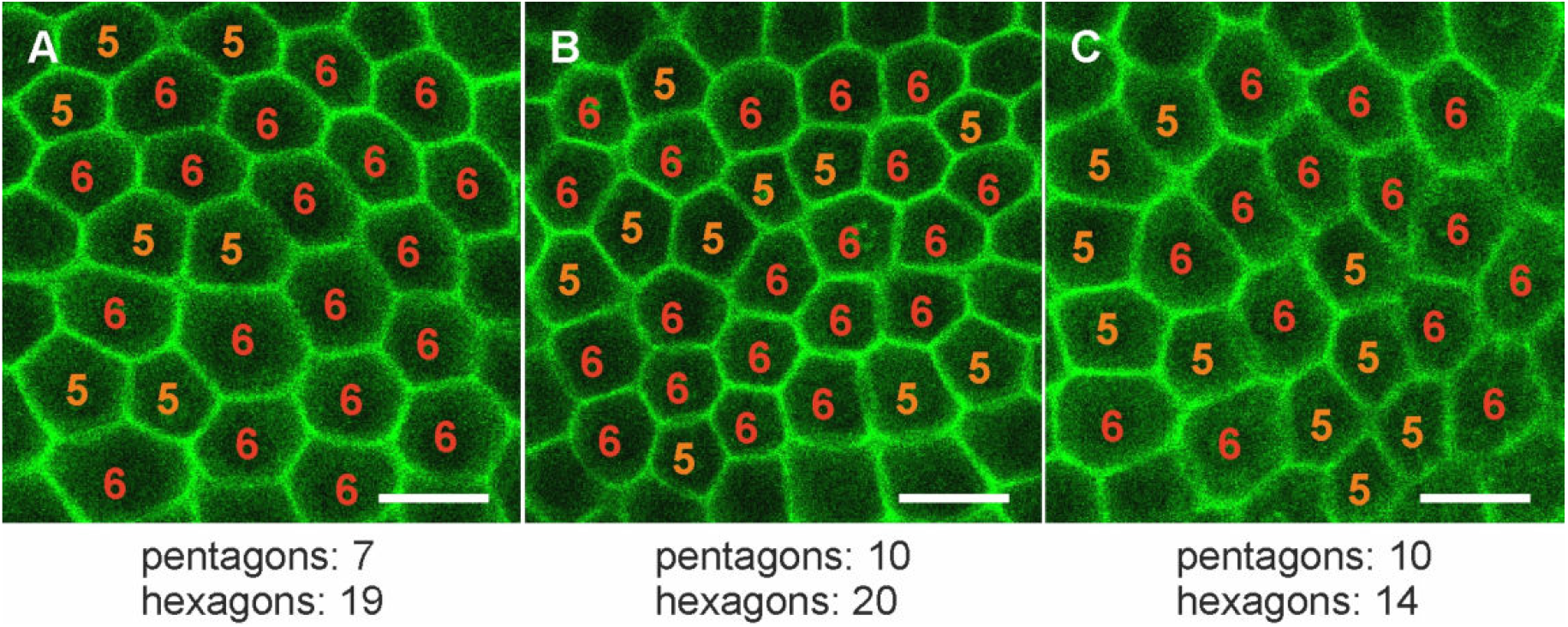
Share of pentagonal and hexagonal somatic CZ3 compartments in middle aged adults (early stage II). Sexually induced transformants expressing the *phII:yfp* gene under the control of the endogenous *phII* promoter were analyzed in vivo for the localization of the PHII:YFP fusion protein. Magnified view of the PhII:YFP-stained compartments surrounding the somatic cells corresponding to CZ3. In areas where no gonidia lie below the somatic cell sheet, the compartments form a pattern of hexagons and pentagons. In the exemplary regions shown here, 27 pentagonal and 53 hexagonal compartments were counted, representing a ratio of roughly 1:2. Scale bars are 20 *µ*m.

**FIG. S5.**
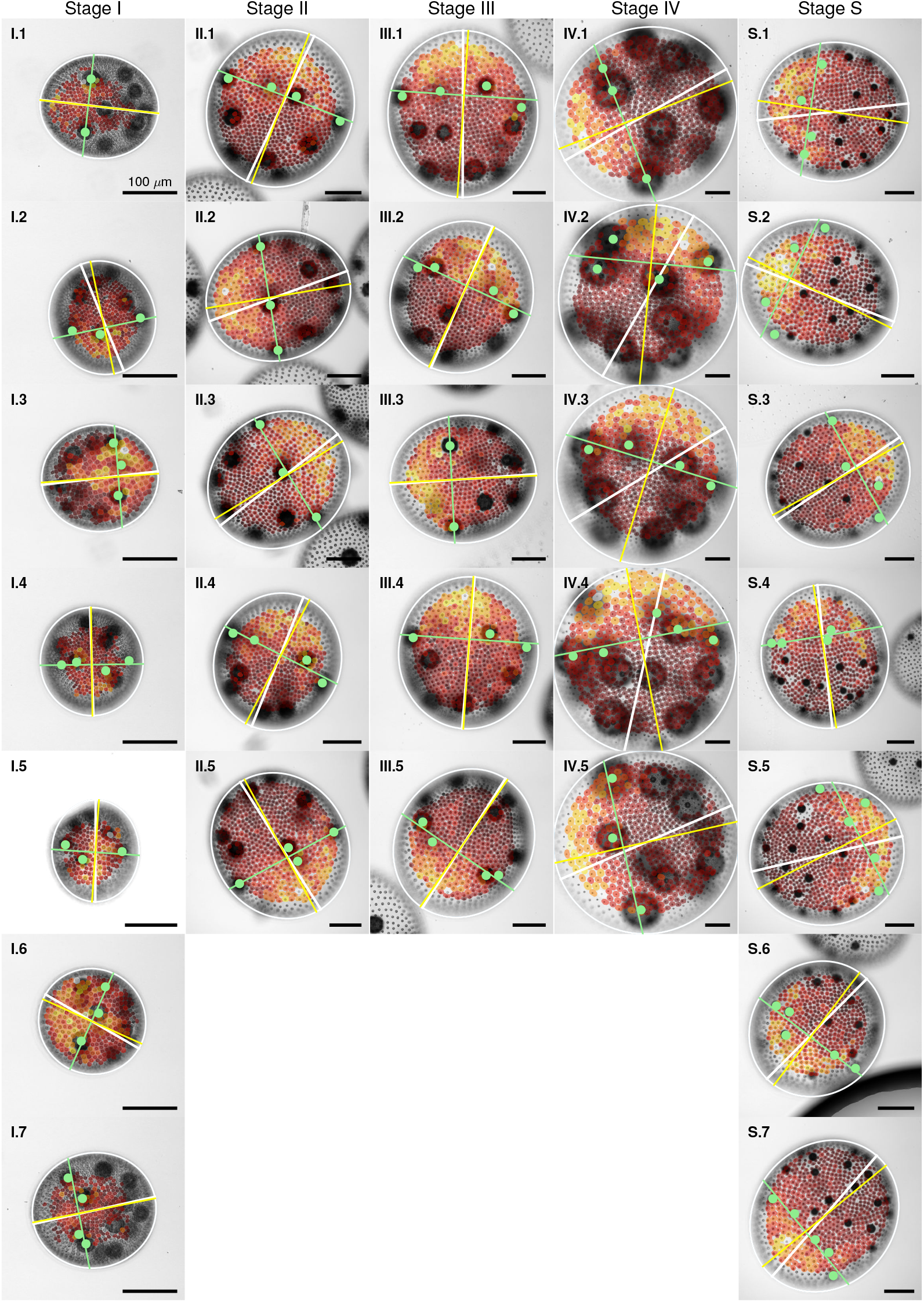
Segmented PhII:YFP signal and anterior-posterior axis identification. Trans-PMT images of spheroid in stage I-IV and S (defined in Fig. 3 in the main document) with posterior-anterior axis (yellow line) estimated as the line passing through the elliptical (white outline) center which is normal to the best-fit line (green) through manually identified offspring (green dots) located in the anterior section. The white line represents the major axis of the ellipsoid. Overlaid are segmentations of the CZ3 compartments (PhII:YFP), colored by area (dark to light by size).

**FIG. S6.**
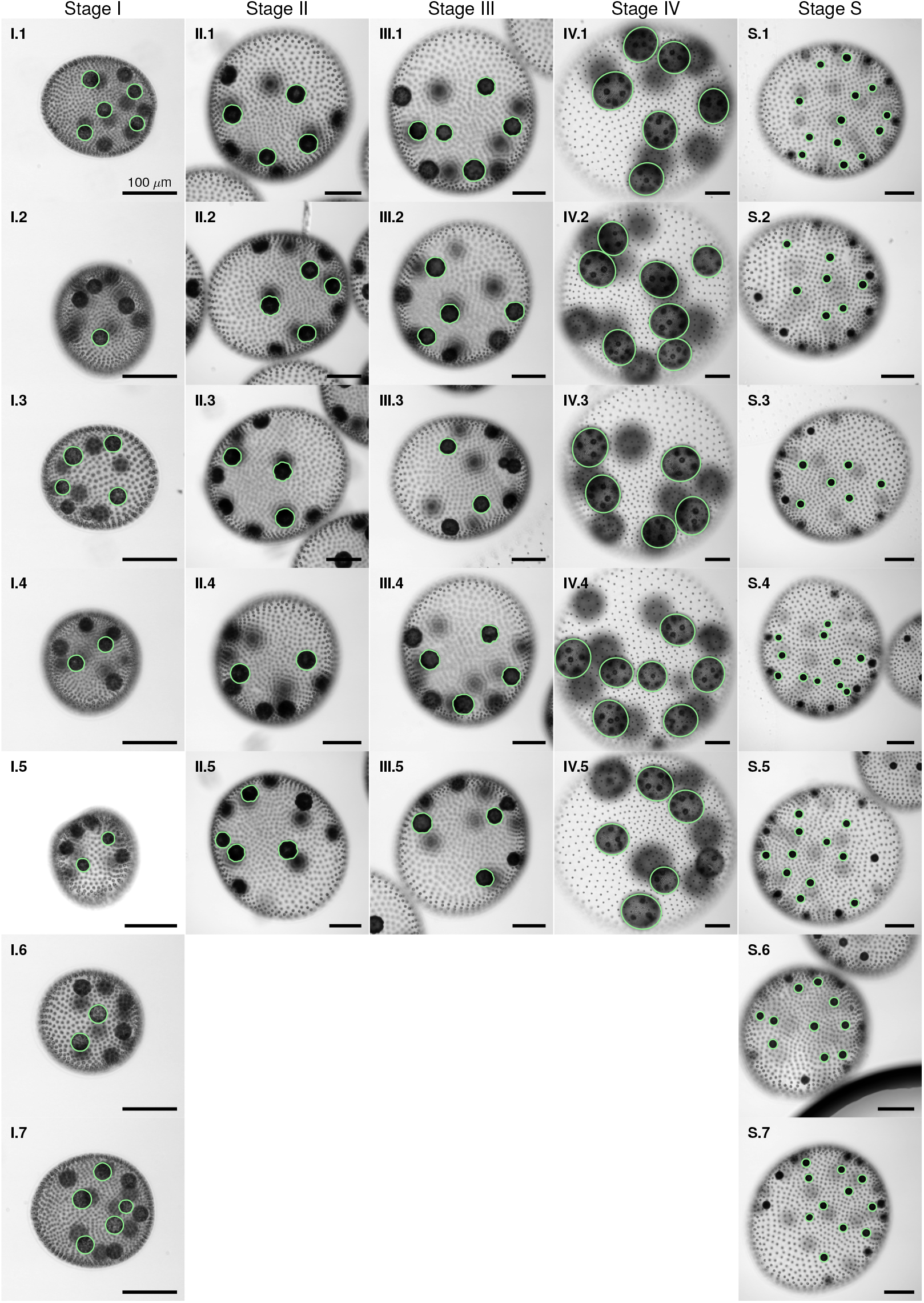
Offspring. Offspring are identified using the same semi-automated procedure as that used for identification of somatic cells and CZ3 compartment geometry. Being far larger than either of those, however, offspring appear in focus at different planes. We identify offspring from planes where clearly in focus, displaying here particular planes for each spheroid. The boundaries are used to estimate the volumetric growth rates displayed in Table 2, main text.

**FIG. S7.**
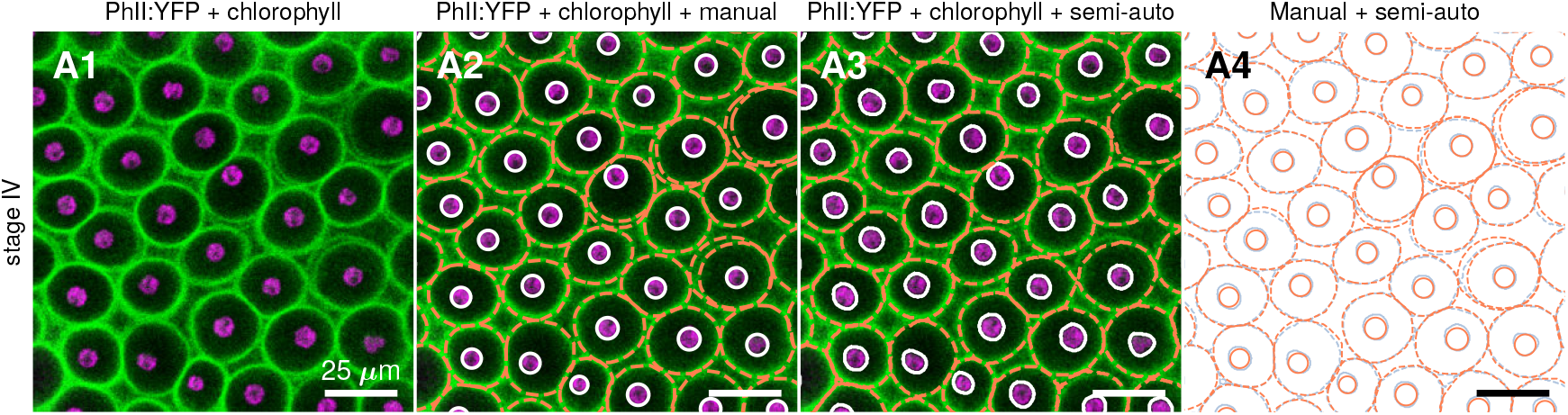
Comparison of semi-automated segmentation against fully manual segmentation. 1. Overlay of YFP fluorescence of PhII:YFP protein (green) and chlorophyll fluorescence (magenta), detected at 650-700 nm. 2. Same as 1 with *manually* segmented CZ3 (orange) and cell boundaries (white), identical to panel A2, Fig. 5 in the main text. 3. Same as 2 with *semi-automatically* segmented (using the procedure described in §2) CZ3 (orange) and cell boundaries (white). 4. Comparison of semi-automated segmentation (blue, CZ3 compartments and cells) with fully manual segmentation (orange, CZ3 compartments and cells).

**FIG. S8.**
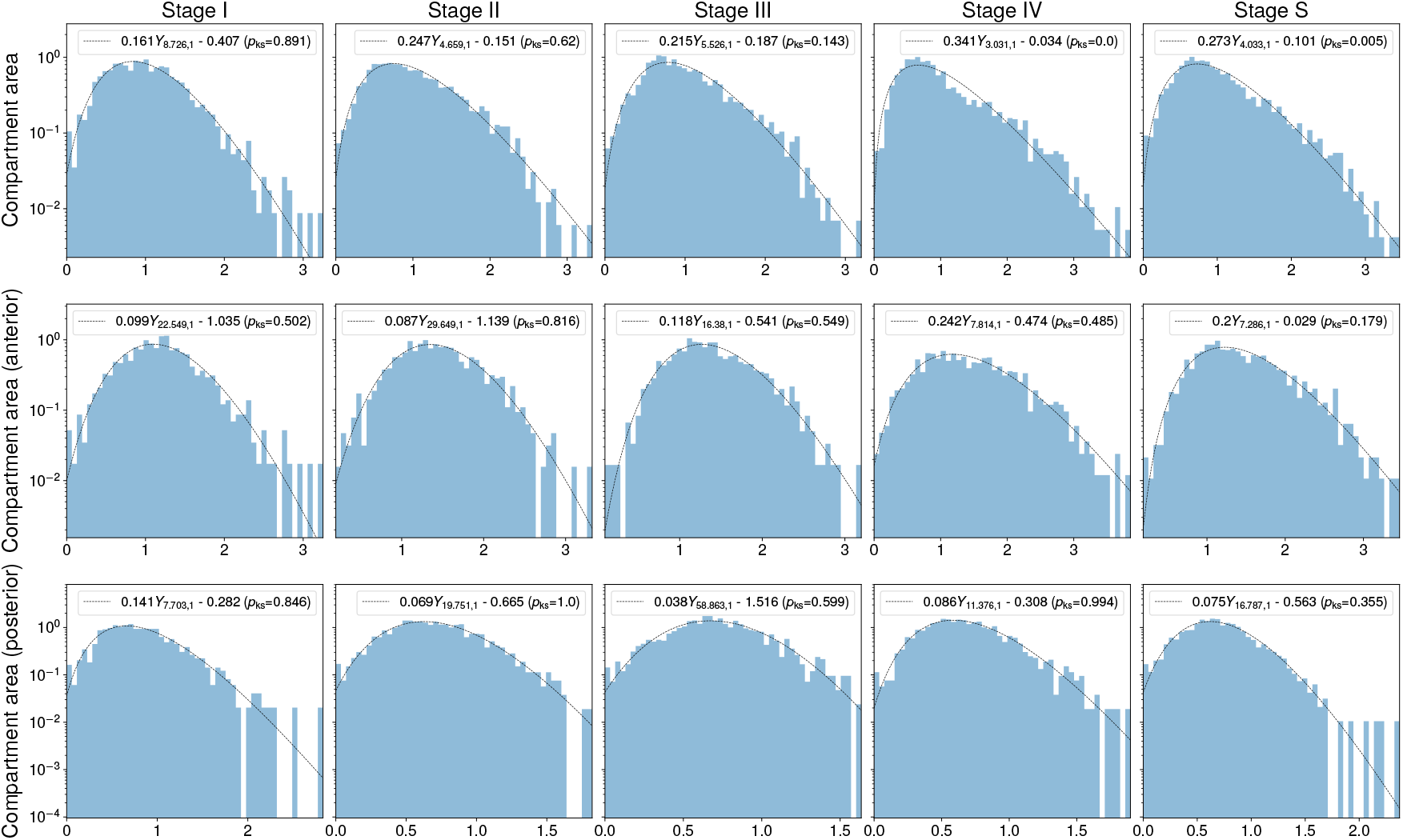
Areas are gamma-distributed throughout growth with changing shape parameters reflecting anteriorposterior differentiation. Areas of CZ3 compartments robustly follow gamma-distributions, as highlighted in Fig. 8B2 and §E, main text. Plotted on y-axes (shared by row) are normalized counts per bin (*n* = 50 bins), and on x-axes are the reduced CZ3 areas ã_cz3_ = (*a*_cz3_ −*a*_min_)*/*(*a*_avg_ −*a*_min_), where *a*_min_ and *a*_avg_ are minimum and average values of *a*_cz3_ for the respective spheroid. Such normalization enables distributional fit across different organisms. The empirical mean is therefore fixed at 1 in these distributions, and the maximum-likelihood fit parameters are *k* and offset of the support (with *λ* determined by the relation between *k* and the fixed mean). A Kolmogorov-Smirnov (KS) goodness-of-fit test is performed and its *p*-value is recorded as *p*_ks_ in each plot. One key observation we make, evident in the first row, is that the extreme degree of anteriorposterior differentiation (shown in Fig. 8B1, main text) suggests that a single fit from this distribution family may not be valid in later stages of the life cycle. Indeed, *p*_ks_ ≈0.9 in stage I, indicating no strong evidence that ã_cz3_ do not arise from a gamma distribution, yet drops below 0.001 by stage IV, supporting rejection of the fit hypothesis in this case. Qualitatively, one observes the formation of “shoulders” in the distribution (exhibiting a loss of log-concavity which does not hold in the gamma distribution for shape parameter *k ≥*1). Separating the data by anterior/posterior hemispheres, however, almost completely resolves the issue and reveals that the respective ã_cz3_ values follow gamma distributions (of very different shape parameters). Lastly, fitted parameters of the sexual stage S place it distributionally closer to Stage IV than to III, which contrasts with the grouping inferred from the spheroid radii (Fig. 10E, main text).

**FIG. S9.**
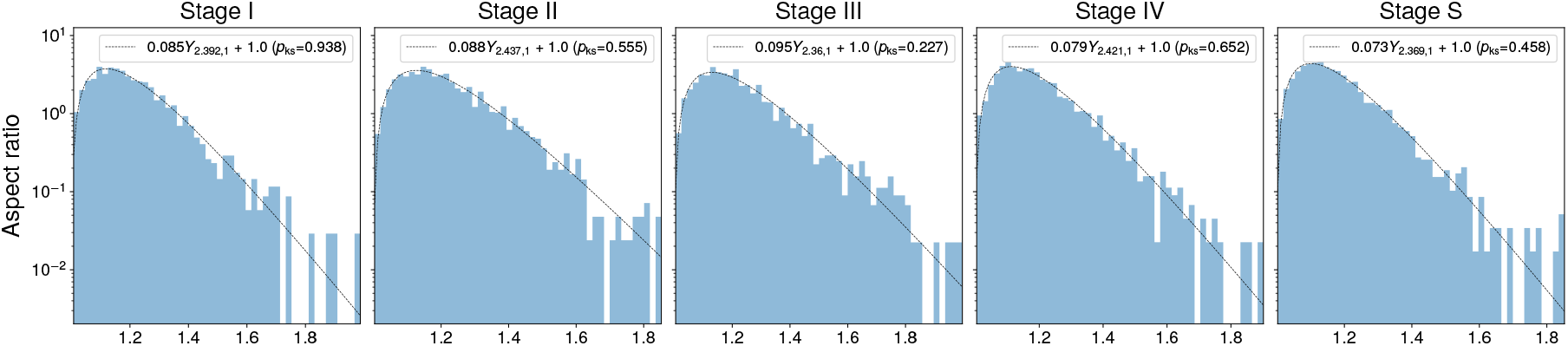
Aspect ratios remain stably gamma-distributed throughout growth. As seen, the *k* parameter remains in a remarkably tight range between 2.35 ™2.45. Maximum-likelihood estimated (MLE) parameters include only rate *λ* and shape *k*, with location fixed at 1. Both this distribution family and the value of the shape parameter for aspect ratios are robustly observed in a variety of living organisms and inert jammed systems [S17]. Kolmogorov-Smirnov tests performed in each case indicate no strong evidence that the data do not follow gamma distributions.

**TABLE S1.** Estimated volumetric growth changes by life cycle stage (I-IV), supplementary to Table 2, main text. Number of juveniles per spheroid were manually estimated from the trans-PMT image (as in Fig. S6), with the average value over all *n* = 29 spheroids being exactly 13. The estimated total offspring volume change (column 3) is defined as this number, times the average estimated juvenile volume per stage (as given in Table 2, main text, computed from segmentation of the trans-PMT image, Fig. S6). Further, assuming a fixed count of 2^11^ ≈2000 somatic cells per spheroid and multiplying this times the average somatic cell volume per stage (again Table 2, main text) yields the estimated total volume change due to somatic cells (column 4). Assuming 2^12^ somatic cells changes the contribution to the overall spheroid volume negligibly, as evident from column 6. Taken together, columns 2, 3, and 4 yield an estimation of the contribution to overall spheroid volume change by offspring, somatic cells, and parental ECM (Table 2, column 5, main text) by subtracting columns 3 and 4 from 2. Lastly, knowledge of the approximate elapsed time between life cycle stages (column 7) yield the corresponding estimated growth rates (columns 8-10 here and Table 2, columns 5-6, main text).

| Stage | Spheroid volume change (est., mm<sup>3</sup>) | Total offspring volume change (est., mm<sup>3</sup>) | Total cell volume change (est., μm<sup>3</sup>) | % spheroid volume change to offspring (est., %) | % spheroid volume change to cells (est., %) | Elapsed time (h) | Spheroid growth rate (est., mm<sup>3</sup>/h) | Total offspring volume growth rate (est., mm<sup>3</sup>/h) | Total cell volume growth rate (est., μm<sup>3</sup>/h) |
| --- | --- | --- | --- | --- | --- | --- | --- | --- | --- |
| I | ↓ | ↓ | ↓ | ↓ | ↓ | ↓ | ↓ | ↓ | ↓ |
| II | 0.040 | 0.0011 | $2.9 \times 10^5$ | 2.8 | 0.73 | 15 | 0.0027 | $7.5 \times 10^{-5}$ | 9.8 |
| III | 0.015 | 0.00019 | $1.9 \times 10^4$ | 1.2 | 0.13 | 6 | 0.0025 | $3.1 \times 10^{-5}$ | 1.6 |
| IV | 0.26 | 0.026 | $9.4 \times 10^4$ | 10 | 0.037 | 16 | 0.016 | 0.0016 | 2.9 |

**TABLE S2.** Summary of changes in mean value along posterior-anterior axis (PA) and by life cycle stage (I-IV, S). Rows 1-5 show the changes in empirical mean value from posterior to anterior end within each life cycle stage. Rows 6-12 show the changes in mean value across successive life cycle stages. While the underlying distributions are highly skewed and mean-based comparisons should be interpreted carefully as noted in §E1 (main text), they provide one quantification of the underlying trends visible in the empirical distributions, Fig. 8, main text.

|  | Cell area | Compartment area | Aspect ratio | Circularity | Offset | Offset (whitened) | Voronoi error |
| --- | --- | --- | --- | --- | --- | --- | --- |
| % change PA in stage I | 14 | 44 | 9.8 | -2.5 | 47 | 23 | 1.7 |
| % change PA in stage II | 13 | 94 | 3.1 | -0.69 | 120 | 65 | 16 |
| % change PA in stage III | 13 | 120 | 1.0 | -0.67 | 82 | 37 | -6.6 |
| % change PA in stage IV | 8.1 | 130 | 14 | -0.53 | 50 | 8.7 | -10 |
| % change PA in stage S | 9.9 | 130 | -1.8 | -1.9 | 200 | 110 | -3.1 |
| Mean value in stage I | 26 μm | 58 μm<sup>2</sup> | 1.2 | 0.88 | 0.54 μm | 0.24 | 0.21 |
| % change stages I to II | 81 | 200 | 0.93 | 1.9 | 140 | 43 | 22 |
| Abs. change stages I to II | 21 | 120 | 0.011 | 0.017 | 0.76 | 0.11 | 0.045 |
| % change stages II to III | 2.4 | 30 | 0.86 | -0.78 | 27 | 12 | -1.4 |
| Abs. change stages II to III | 1.1 | 52 | 0.010 | -0.070 | 0.35 | 0.042 | -0.036 |
| % change stages III to IV | 12 | 170 | -2.7 | 1.8 | 53 | -5.6 | 16 |
| Abs. change stages III to IV | 6.0 | 370 | -0.34 | 0.016 | 0.87 | -0.22 | 0.039 |

**FIG. S10.**
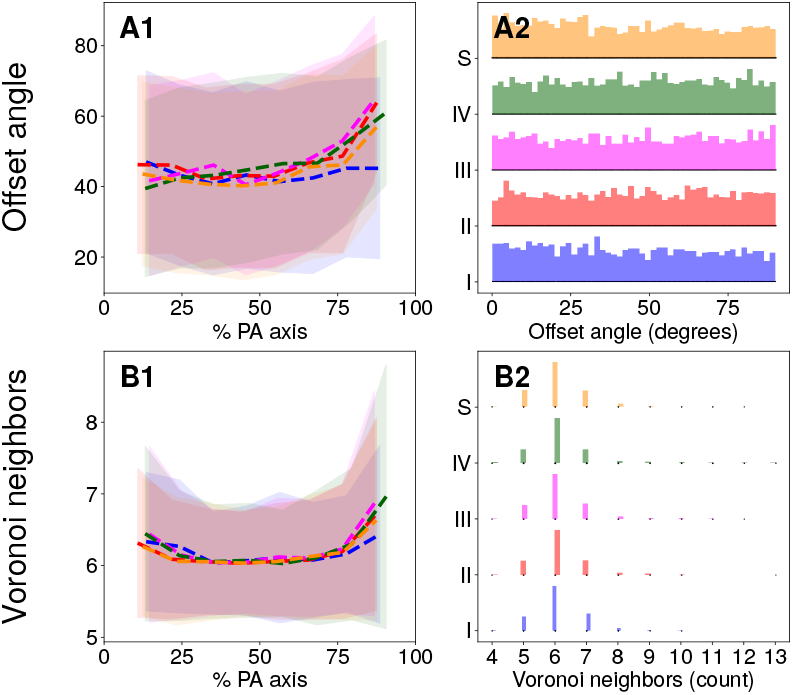
Additional invariances in the stochastic geometry of the CZ3 compartments across the life cycle. Rows A and B follow the same formatting as Fig. 8, main text. Row A shows variation in the offset angle *θ*_cell_ = arccos(*Δ****x***· ***v***| */*(∥*Δ****x***∥∥ ***v***∥)) between the somatic cell offset vector *Δ****x*** (Table 1, main text) and the principal stretch axis ***v*** of the CZ3 compartment (given by the eigenvector corresponding to *λ*_max_). Since the latter has no polarity, *θ*_cell_∈ [0, 90] degrees. Panel A1 shows that there is little variation along the PA axis or by life cycle, with the exception of upward tails toward the anterior pole in stages II-IV and S. Panel A2 reveals furthermore that *θ*_cell_ effectively follows a uniform distribution over its support throughout the lifcycle, indicating that there is no correlation between the somatic cell offset vector *Δ****x*** and the principal stretch axis. This is perhaps surprising given that one might naively expect translation of the cell along the deformation axis of its compartment. Row B shows, in the same format, the number of neighbors (edges) of each Voronoi partition. Panel B1 again shows that there is almost no variation along the PA axis or by life cycle stage, providing quantitative evidence that the topology of the CZ3 space partition does not change during growth. Panel B2 displays the distribution of Voronoi neighbors by life cycle stage, whose mean is around 6, consistent with Euler’s theorem for Voronoi tessellations of the sphere (§2 D).

**FIG. S11.**
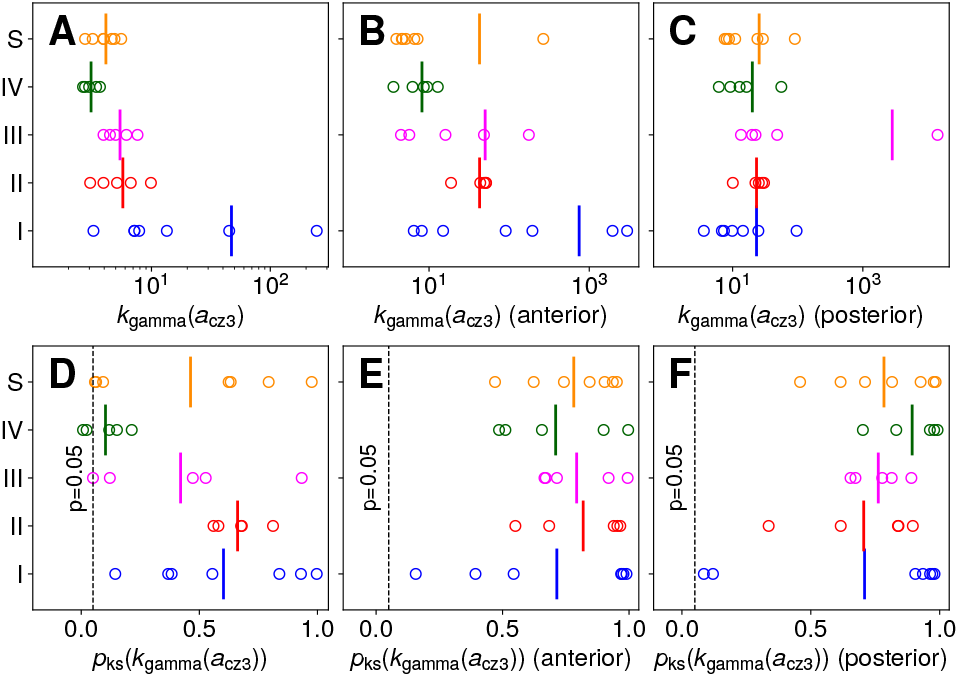
Increasing area polydispersity during the life cycle is primarily due to changes in the anterior hemisphere. Panel A is the same as Fig. 10B (main text) while panels B and C show the same metrics separated by anterior and posterior regions respectively. As seen, B shows a decrease in *k* (in increase in disorderedness as explained in main text) of the anterior configuration by life cycle stage, while C shows relatively stable values in the posterior configuration. The former therefore likely underlies the increasing disorderedness observed in the whole spheroid (panel A). Panels D-F display the respective *p*-values (*p*_ks_) of a Kolmogorov-Smirnov goodness-of-fit test performed in each instance of the *k*_gamma_ value reported in panels A-C. All panels D-F show that stage I can exhibit varying degrees of goodness-of-fit, which is likely reflected by the large spread of *k*-values in stage I (panels A-C). However, in the progression from stages II-IV and S, *p*_ks_ drops significantly, sometimes below the threshold (plotted as dashed vertical lines), as expected from Fig. S8, despite the qualitatively good match observed in row 1 of that figure. As in Fig. S8 rows 2-3, this is remedied by separating into anterior/posterior hemispheres, at which point the data collapses well, indicated by the sharp increases in *p*_ks_ in panels E-F. Remarkably, this increase in goodness-of-fit accompanies a clear observation in panels B-C that the anterior hemisphere is primarily responsible for increasing disorder in the somatic CZ3 configuration.

## Notes

http://doi.org/10.5281/zenodo.14066435

### Competing Interest Statement

The authors have declared no competing interest.

